# Distinct organization of two cortico-cortical feedback pathways

**DOI:** 10.1101/2020.02.27.968792

**Authors:** Shan Shen, Xiaolong Jiang, Federico Scala, Jiakun Fu, Paul Fahey, Dimitry Kobak, Zhenghuan Tan, Jacob Reimer, Fabian Sinz, Andreas S. Tolias

**Affiliations:** Center for Neuroscience and Artificial Intelligence, Baylor College of Medicine, Houston, Texas, USA; Department of Neuroscience, Baylor College of Medicine, Houston, Texas, USA; Jan and Dan Duncan Neurological Research Institute at Texas Children’s Hospital, Houston, Texas, USA; Institute for Ophthalmic Research, University of Tübingen, Germany; Bernstein Center for Computational Neuroscience, University of Tübingen, Tübingen, Germany; Institute for Bioinformatics and Medical Informatics, University of Tübingen, Tübingen, Germany; Department of Electrical and Computational Engineering, Rice University, Houston, Texas, USA

## Abstract

Neocortical feedback is critical for processes like attention, prediction, and learning. A mechanistic understanding of its function requires deciphering its cell-type wiring logic. Recent studies revealed a disinhibitory circuit between motor and sensory areas in mice, where feedback preferentially targets vasointestinal peptide-expressing interneurons, in addition to pyramidal cells. It is unknown whether this circuit motif is a general cortico-cortical feedback organizing principle. Combining multiple simultaneous whole-cell recordings with optogenetics we found that in contrast to this wiring rule, feedback between the hierarchically organized visual areas (lateral-medial to V1) preferentially activated somatostatin-expressing interneurons. Functionally, both feedback circuits temporally sharpened feed-forward excitation by eliciting a transient increase followed by a prolonged decrease in pyramidal firing rate under sustained feed-forward input. However, under feed-forward transient input, the motor-sensory feedback facilitated pyramidal cell bursting while visual feedback increased spike time precision. Our findings argue for multiple feedback motifs implementing different dynamic non-linear operations.

## Introduction

The mammalian neocortex is composed of hierarchically organized areas, where cortical areas are reciprocally connected with both feed-forward and top-down feedback long-range projections. Feedback connections are believed to participate in many important brain functions, such as attention ^1–3^, prediction ^4–8^, and shaping activity based on context ^9–13^. Despite the importance of cortical feedback, its precise cell-type wiring logic is still not fully understood, preventing a mechanistic understanding of its modulatory role in cortical processing.

The mammalian sensory neocortex is organized in a six-layer structure, composed of distinct cell types wired in canonical circuit motifs. For example, inhibitory interneurons in cortical circuits are highly heterogeneous grouped into transcriptomic and morphological cell classes, hypothesized to exert distinct functional roles ^14–17^. Therefore, it is critical to deciphering the cell-type-specific wiring of feedback connections. Among GABAergic interneurons, parvalbumin (PV), somatostatin (SOM), and vasointestinal peptide (VIP) expressing interneurons are the three main non-overlapping cell classes that comprise more than 80% of the GABAergic interneurons ^15^. Importantly, these cell classes follow specific rules governing their connections with other interneuronal types and local excitatory neurons ^14,18,19^. Specifically, a key connectivity rule is that VIP+ interneurons preferentially inhibit SOM+ interneurons which in turn inhibit local pyramidal cells ^14,18,20^. Recent studies indicate that feedback projections primarily recruit this disinhibitory disynaptic circuit by preferentially activating VIP+ interneurons ^21–24^, supporting the view that the primary effect of feedback is a disinhibition of target areas ^25^. However, the cortical feedback projections studied so far mainly focused on connections between brain regions across different modalities such as between motor and sensory areas ^20,22,24^. Therefore, it is not clear if feedback projections recruit the same disinhibitory circuit between hierarchically organized areas within the same sensory modality and if the feedback-VIP+ circuit motif described so far is a universal organizing principle of cortical feedback

The mouse visual cortex is an ideal model to address this question given its hierarchically organized extrastriate areas ^26–34^. Here we focused on feedback connections from the lateral-medial area (LM), which is believed to be analogous to area V2 in primates ^31^, to area V1. We compared the organization of this within-hierarchy feedback to the previously-studied feedback pathways across modalities, from the vibrissal primary motor cortex (vM1) to vibrissal S1 (vS1), to examine if there is a general rule governing the organization of feedback pathways across these two different pathways. We found major differences in both the cell-type-specific wiring rules, distribution of feedback projecting axons in terms of cortical areas and layer specificity and the functional impact of feedback projections between these two pathways. LM to V1 feedback connected more strongly to SOM+ cells than to VIP+ cells while vM1 to vS1 feedback showed the opposite pattern, consistent with previous studies ^21^. When paired with sustained positive current injection into the cell bodies of pyramidal cells in either V1 or vS1, feedback projections had a similar effect on the activity of pyramidal cells temporally sharpening the feed-forward excitation by eliciting a transient increase followed by a sustained decrease in firing rate. However, when paired with a brief positive feed-forward current pulse, vM1 to vS1 feedback facilitated bursting of Layer 5 (L5) intrinsically bursty (IB) cells. In contrast, under the same brief feed-forward input, LM to V1 feedback increased the probability of a second spike but eliminated the rest of spikes, in agreement with temporal sharpening. Our results argue for multiple feedback circuit motifs specialized for distinct dynamic non-linear operations.

## Results

### Distribution of feedback axon terminals

To study the projection pattern and connectivity of feedback pathways, we injected the adeno-associated virus (AAV2/1) expressing channelrhodopsin-2 (ChR2)-YFP in either LM or vM1 (Figure 1A, Methods). LM was identified with intrinsic optical imaging ^26,35^; Figure 1B), while vM1 was identified stereotaxically (0.9 mm lateral and 1.1 mm anterior of bregma; ^21,36^; Figure 1A, see Methods).

**Figure 1:**
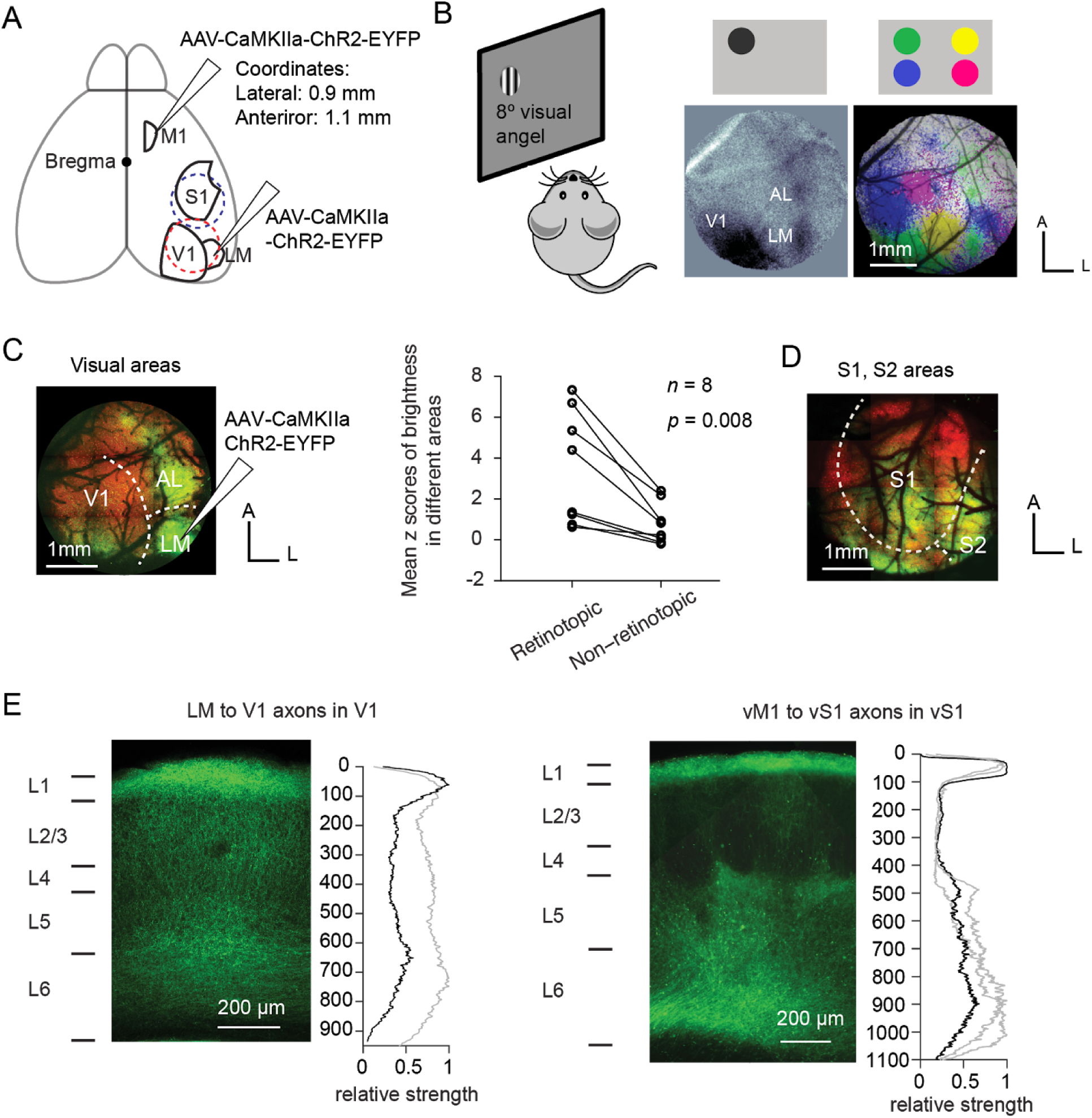
Distribution of feedback axon terminals of the two feedback pathways. **(A)** Anatomy of the areas V1, LM, vS1, and vM1. The virus was injected in either vM1 or LM in different animals. Craniotomies were made to expose the visual cortical areas (red dashed circle) or somatosensory cortical areas (blue dashed circle) to examine the expression of ChR2-YFP in the axon terminals. **(B)** Intrinsic imaging to identify visual areas V1 and LM. Left: experimental paradigm. Grating stimuli drifting in horizontal or vertical directions were shown to the left eye of the animal, on one of the four locations (top lateral, top medial, bottom lateral, bottom medial) on the monitor. A CCD camera was used to record the intrinsic autofluorescence of the brain from a craniotomy on the right hemisphere, exposing the visual areas. Middle: intrinsic imaging map of the stimulus in the top lateral corner. Note that AL and LM were well separable. Right: intrinsic imaging map of all stimuli in four locations on the screen. Different colors represent brain areas that were responsive to stimuli in different locations (green: top lateral; yellow: top medial; blue: bottom lateral; magenta: bottom medial). Scale bar: 1 mm. **(C)** LM to V1 feedback projections target the retinotopic corresponding area in V1. Left: virus expression of the same animal in (B). The virus was injected in the “green location” (cells responsive to top lateral stimulus, refer to panel B) in LM, and the axon terminals mainly targeted the “green location” in V1. Right: mean z scores of the fluorescence for both retinotopically corresponding areas and non-corresponding areas in V1, for 8 animals. Fluorescence strength of the retinotopic area was significantly higher than that of the non-retinotopic area (*p* = 0.008, *n* = 8 animals, Wilcoxon signed-rank test). **(D)** Dorsal view of vM1 to vS1 feedback projections. Axon terminals were widely distributed in the somatosensory areas. **(E)** Laminar distribution of LM to V1 (left) or vM1 to vS1 (right) axon terminals. Left: the fluorescent image of a coronal slice in V1 or vS1. Right: Relative strength of fluorescence (averaged over 100 μm horizontally) as a function of depth, normalized to the maximum. The black line indicates the distribution of the axon terminals corresponding to the image on the left, and the gray lines indicates the distribution of other slices.

Two to four weeks later, we used two-photon microscopy to image the fluorescence within a 3 mm cranial window centered on either V1 (centered at 3.0 mm lateral, 1.5 mm anterior to lambda, Figure 1A, red dashed circle, and Figure 1C) or vS1 (centered at 3.5 mm lateral and 1.5 mm posterior to bregma, Figure 1A, blue dashed circle, and Figure 1D). Consistent with a recent study ^37^, feedback projections from LM mainly targeted the retinotopically-matched area in V1 (Figure 1C and D; z-score of fluorescence in retinotopically-matched and unmatched areas in V1; *p* = 0.008, Wilcoxon signed-rank test, *n* = 8 animals), confirming a spatial specificity in the LM to V1 feedback projections ^37^. On the other hand, vM1 to vS1 projections broadly covered vS1 and other areas such as secondary somatosensory area and posterior parietal cortex (Figure 1D).

Coronal slices revealed that LM to V1 axon terminals spanned all layers, with L4 having the sparsest projections (Figure 1E, left). In contrast, vM1 to vS1 axon terminals were concentrated in L1 and deep layers (L5 and L6), and were very sparse in L2/3 and L4 (Figure 1E, right), consistent with previous reports ^36^.

### Cell-type wiring logic of the two feedback pathways

To identify the cellular targets of both feedback pathways, we performed multi-cell simultaneous whole-cell recordings on acute brain slices prepared from either V1 or vS1 areas (Figure 2A, see Methods). We recorded from excitatory cells and the major genetically identified classes of interneurons in Layers 1, 2/3, 4, and 5 (*n* = 503 cells in total). Distinct interneuronal classes were identified using different mouse lines (PV-Cre/Ai9, SOM-Cre/Ai9, or VIP-Cre/Ai9) and were further confirmed by *posthoc* analyses of their morphological and electrophysiological properties (See Methods, Supplementary Figure S1, S2). We recorded excitatory postsynaptic currents (EPSCs) and excitatory postsynaptic potentials (EPSPs) from each neuron evoked by photostimulation of ChR2-YFP-expressed in the axon terminals of feedback projections from LM or vM1 (2 ms; 470 nm, Figure 2B-E for EPSP examples).

**Figure 2.**
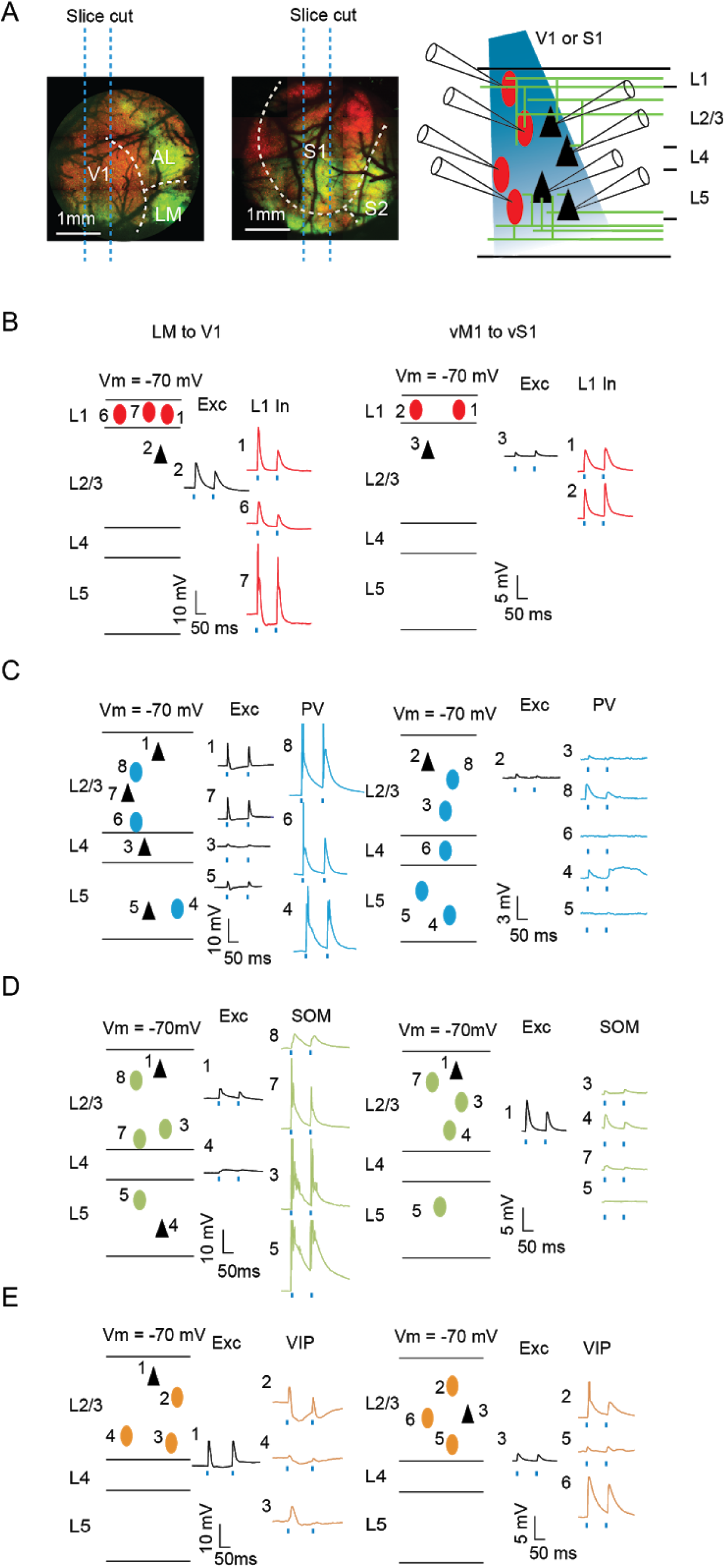
Example slices for connectivity. **(A)** Experimental paradigm. Left: Parasagittal slices containing V1 or vS1 Right: LM or vM1 were excluded from the slices, while the axon terminals from LM or vM1 pertained in the slices. **(B)** Example slice recording of L1 interneurons (red ovals) and pyramidal cells (black triangles) in either V1 (left) or vS1 (right). Numbers refer to the channel number of the recording system. **(C)** Example slice recording of PV+ interneurons (blue ovals) and pyramidal cells (black triangles) in either V1 (left) or vS1 (right). **(D)** Example slice recording of SOM+ interneurons (green ovals) and pyramidal cells (black triangles) in either V1 (left) or vS1 (right) **(E)** Example slice recording of L2/3 VIP+ interneurons (orange ovals) and pyramidal cells (black triangles) in either V1 (left) or vS1 (right).

In both V1 and vS1, we found a high proportion of cells responsive to feedback stimulation across most cortical layers and cell classes (overall 85.7%, 431/503), except for L4 in vS1, where a very low number of excitatory cells (1/13), none of the SOM+ cells (0/8) and only about half (8/14) of the PV+ cells were responsive (Figure 3A, see Methods for detailed criteria for “responsiveness”).

**Figure 3.**
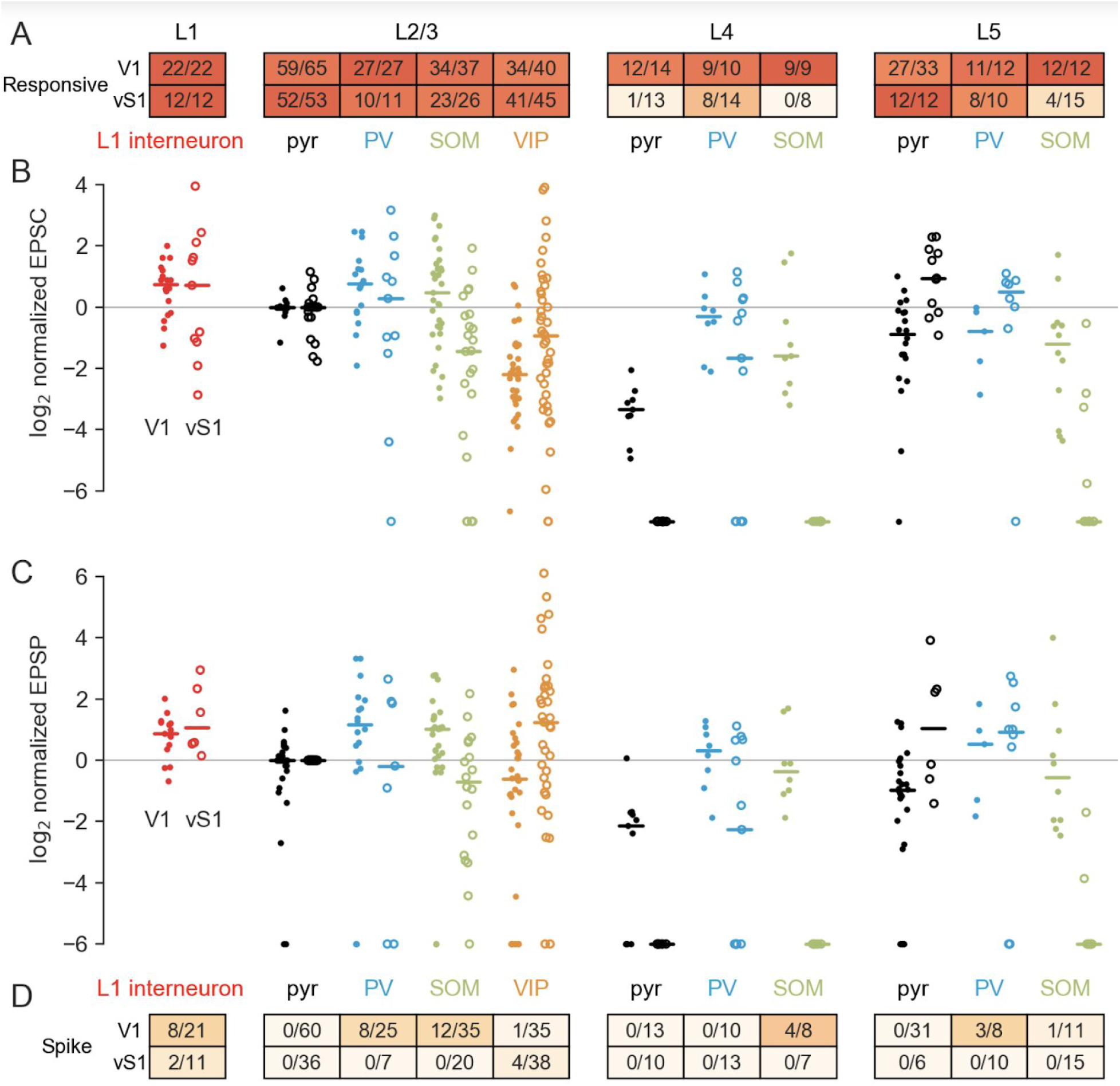
Summary of activities in V1 and vS1 in response to feedback excitation. **(A)** The proportion of responsive cells in V1 or vS1 (number of responsive cells/total number of recorded cells). The color in the table indicates the probability level, same for panel C. **(B)** The log2 EPSC of cells in either V1 (solid dots) or vS1 (open dots) normalized to the average activity of L2/3 pyramidal cells. **(C)** Same as (B), for normalized EPSP. **(D)** The proportion of spiking cells in V1 or vS1 (number of spiking cells/total cells recorded in the current-clamp mode).

We performed a subset of experiments in the presence of tetrodotoxin (TTX) and 4-aminopyridine (4-AP) and confirmed that the majority of the evoked events were monosynaptic (Methods, Supplementary Figure S3). In agreement with that, we never found a pyramidal cell in any layer in either area firing in response to the feedback activation alone (Figure 3D). Since poly-synaptic events could only be elicited if local excitatory neurons are driven to fire, these results also suggest that most of the recorded feedback-evoked events were monosynaptic.

Although most of the recorded cells were responsive to feedback activation, the strengths of connections varied considerably, across both the cell classes and feedback pathways (Figure 2B-D). To compare the strength of responses to feedback across layers and cell types recorded in different brain slices and animals, we normalized the amplitudes of EPSCs and EPSPs to the mean amplitudes of the L2/3 pyramidal cells recorded from each slice, similar to previous approaches ^21,24^, and report the log2 normalized amplitudes (Figure 3B and C). We found that for both feedback pathways, the normalized EPSPs and EPSCs varied considerably across different cell classes (Figure 3B and C, *p* < 10^−6^ for each of the four separate Kruskal-Wallis tests for EPSPs and EPSCs in V1 and vS1).

To quantify these differences, we performed statistical comparisons between normalized EPSPs and EPSCs in all pairs of neural types within each feedback pathway, and also compared the responses of the same neural type across the two pathways (we used Connover’s *posthoc* tests, permutation tests, and Mann-Whitney-Wilcoxon tests as appropriate, see Methods for details). Given a large number of statistical tests (55×4 + 10×2 = 240 overall), we report all *p*-values with Benjamini-Hochberg adjustment that allows control of the false discovery rate (Supplementary Tables S1-S4). We found that some groups of neurons in both feedback pathways had consistently higher feedback responses than L2/3 pyramidal cells (Figure 3B). For example, the median log2 normalized EPSPs of L1 interneurons were 0.88 (V1) and 1.1 (vS1; Figure 3C, Table S1-S4 for statistical tests), consistent with the concentrated feedback axon terminals present in L1. At the same time, some other groups of neurons had consistently lower responses than L2/3 pyramidal cells. For example, in V1, the median log2 normalized EPSC and EPSP of L4 excitatory cells were -1.7 and -2.3, respectively (Figure 3B and C), while in vS1, nearly all L4 excitatory cells were not responsive to vM1 feedback activation (Figure 3A-C). These results were consistent with sparse axon terminals present in L4 (Figure 1C and D).

A prominent difference between the two feedback pathways was in the responses of SOM+ neurons. In V1, the median log2 normalized EPSCs and EPSPs of SOM+ neurons were 0.47 and 1.0 for L2/3, -1.6 and -0.36 for L4, and -1.2 and -0.55 for L5, respectively, all of which were higher than their counterparts in vS1. In vS1, the median log2 normalized EPSCs and EPSPs for L2/3 SOM+ neurons were -1.4 and -0.71 (Figure 3B and C) and the SOM+ neurons in L4 and L5 were hardly responsive (Figure 3A). Moreover, vM1 feedback never elicited spikes in any of the SOM+ neurons in vS1 (0/48 across the three layers, Figure 3D). In contrast, in V1, feedback elicited spiking activity in a substantial fraction of SOM+ neurons (17/54).

Another prominent difference between the two feedback pathways was in the responses of VIP+ neurons in L2/3. In V1, the median log2 normalized EPSC and EPSP of L2/3 VIP+ cells were -2.2 and -0.62 (Figure 3B and C), which were lower than their counterparts in vS1, with median log2 normalized EPSC and EPSP to be -0.93 and 1.2 (Figure 3B and C). Notably, other than L1 interneurons, VIP+ neurons were the only neurons in vS1 that spikes were elicited by feedback (4/38, compared to 0 spikes in all other neural types, Figure 3C). In contrast, in V1, spikes were elicited from a lower proportion of VIP+ cells (1/35) than other classes of interneurons, such as SOM+ and PV+ interneurons (Figure 3D).

In summary, we found that both feedback pathways targeted pyramidal cells in L2/3 and L5, L1 interneurons, and PV+ interneurons in L2/3, L4 and L5. Consistent with previous reports ^21^, vM1 to vS1 feedback also strongly targeted L2/3 VIP+ cells but had weaker connections to L2/3 SOM+ cells, and very few connections to SOM+ cells in other layers. In contrast, here we found that LM to V1 feedback strongly targeted L2/3 SOM+ cells but had significantly weaker input to L2/3 VIP+ cells. In addition, excitatory feedback responses in L4 were much weaker in vS1 than in V1 (Figure 3B and C).

The above analysis was performed using L2/3 excitatory responses as the baseline to account for inter-animal and inter-slice variability (e.g. due to the different amount and titer of the injected virus). The non-normalized (raw) responses of the L2/3 excitatory cells were lower in vS1 than in V1 (Supplementary Figure S4). Since on average we injected a similar amount of virus into both areas, this might reflect a difference in the overall connectivity strength between the two feedback pathways. This possible difference, however, does not affect our main conclusions. The raw responses of SOM+ and VIP+ neurons showed the same pattern as the normalized responses (Supplementary Figure S4).

### Feedback projections in both pathways temporally sharpen the firing pattern of V1/vS1 principal cells, when combined with sustained feed-forward input

Next we investigated the function of feedback projections by studying how they modulate the activity of recipient excitatory neurons in V1 or vS1, which are the principal output cells representing visual or tactile information. We first examined how feedback activation interacted with tonic depolarizing current injection designed to mimic sustained feed-forward input.

We injected 300 ms positive current steps in whole-cell configuration to the cell bodies of principal cells in layers 2-5 that drove trains of action potentials in these cells. We paired the depolarizing current injection with a 20 ms light pulse to activate the feedback axon terminals (Figure 4A), randomizing the timing of the light stimulation between 100 and 200 ms after the onset of the current injection. LED-off trials were interleaved with LED-on trials (Figure 4A).

**Figure 4.**
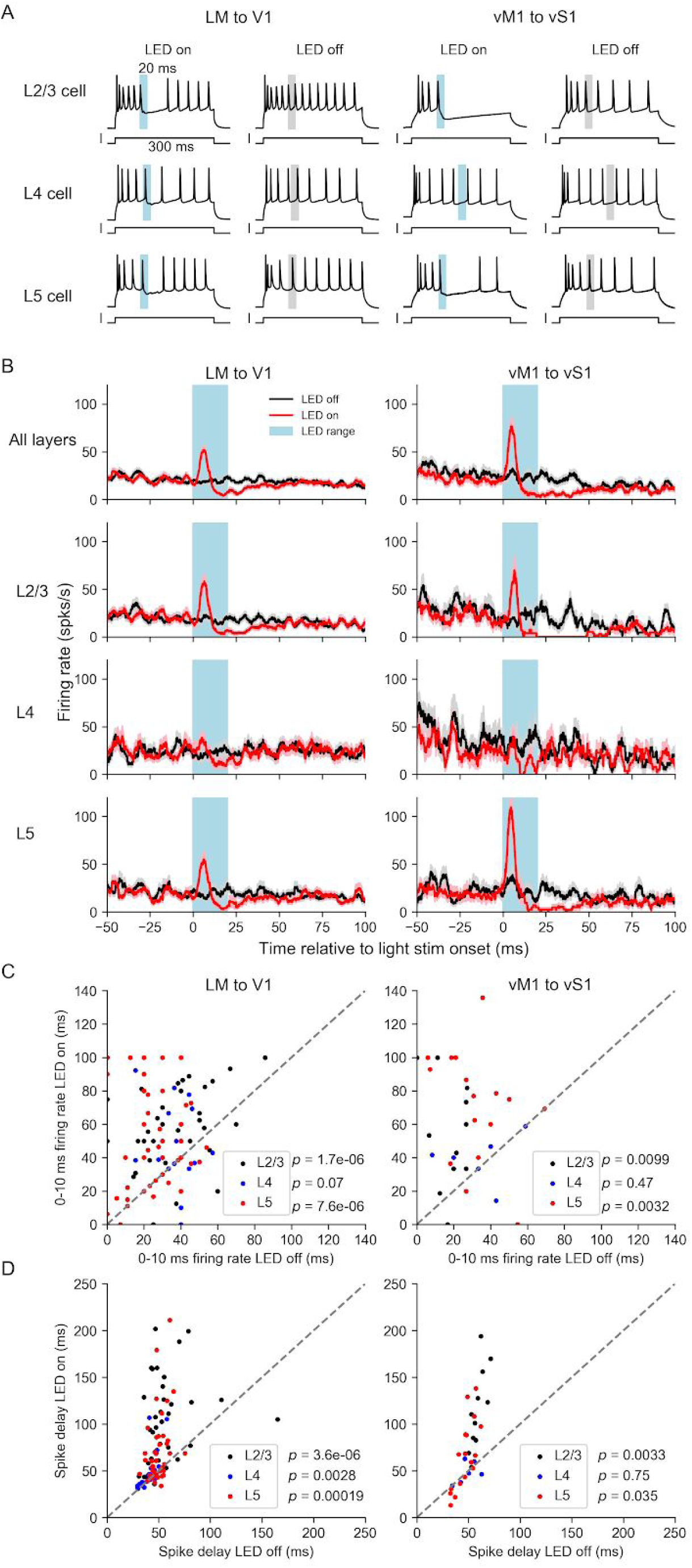
Feedback activity temporally sharpens the firing patterns of pyramidal cells in V1 or vS1 in both feedback pathways. **(A)** Examples of feedback modulation on the firing patterns of pyramidal cells in L2/3 (top), L4 (middle), and L5 (bottom) of V1 (left) or vS1 (right). Cells were driven to fire with sustained positive current injection. In LED-on trials (left plots), 20 ms LED stimulus (blue bar) was delivered. LED-off trials (right plots) were paired with the LED-on trials. Gray bars mark the LED stimulus range of the corresponding LED-on trials. **(B)** Peristimulus time histogram of pyramidal cells for feedback on trials (red, solid line: mean, shade: s.e.m across cells, same for later) and feedback off trials (black). **(C)** Firing-rate in feedback off trials vs feedback on trials within the time range of 0 to 10 ms after the LED onset for pyramidal cells in L2/3 (black dots), L4 (blue dots) and L5 (red dots). **(D)** Time delay relative to the LED onset of the first spike after LED stimulus time range (20 ms) in feedback-on trials vs feedback-off trials for pyramidal cells in L2/3 (black dots), L4 (blue dots) and L5 (red dots).

These experiments revealed consistent effects of feedback modulation in L2/3 and L5 for both feedback pathways: LED stimulation reliably elicited a spike within 10 ms after the LED onset, and delayed subsequent spikes even beyond the 20 ms illumination period (Figure 4A, for more examples, refer to Supplementary Figures S6 and S7). Compared to LED-off trials, the firing rate of neurons during LED-on trials displayed a sharp peak within 10 ms after the LED onset followed by a decrease below the baseline due to the delay of subsequent spiking (Figure 4B). To quantify the effect during the excitation, we compared the firing rate of LED-on and -off trials within 10 ms relative to the LED onset (Figure 4C). To quantify the subsequent delay in spiking after the feedback stimulation, we examined the timing of spikes after stimulation offset (or the matching time points in the LED-off trials). Optogenetic feedback stimulation caused a 2 to 4 fold increase in initial firing (within the first 10 ms following LED-on) in both L2/3 and L5 in both V1 and vS1 (Figure 4C, Wilcoxon signed-rank test, *p* <= 0.0099, *n* >= 11 neurons for all four comparisons), as well as an increased delay of 1.2 to 2 fold in the appearance of the first spike following stimulation offset (Figure 4D, *p* <= 0.035, *n* >= 11 neurons for all four comparisons). Following the terminology of previous work on a different circuit ^38^, we call this effect “temporal sharpening”. This effect was weaker in L4 neurons in V1, with a non-significant increase in the initial firing rate (Figure 4C left, *p* = 0.07, *n* = 20 neurons) and a significant decrease in the delay of the subsequent spikes (Figure 4D left, *p* = 0.0028, *n* = 20 neurons). However, there was no effect in L4 neurons in vS1 (Figure 4C and D right), consistent with the low connection strength to L4 neurons.

This temporal sharpening effect in pyramidal cells in L2/3 and L5 was consistent with the connectivity profile of the feedback circuits, where feedback projections in both pathways connected to both excitatory neurons and inhibitory interneurons (Figure 3). Direct connection from feedback to pyramidal cells potentially increases the firing probability right after the onset of the feedback activation, while disynaptic inhibition (PV+ cells and SOM+ cells in V1, and PV+ cells in vS1) likely mediates the delay in subsequent spiking. In summary, for both LM to V1 and vM1 to vS1 feedback activation temporally sharpened sustained feed-forward excitation by eliciting a transient increase followed by a prolonged decrease in the firing rate of pyramidal cells in L2/3 and L5.

### Only vM1 to vS1 feedback facilitates bursting of intrinsically bursty cells when paired with brief feed-forward input

We next examined the regulatory effect of the feedback activations when paired with brief, temporally-precise feed-forward input. Since previous studies have shown that feedback projections synapse on apical dendrites ^39^ and the activity on apical dendrites facilitates bursting in L5 intrinsically bursty (IB) neurons in S1 ^40^, we hypothesized that feedback will regulate the burstiness of L5 IB neurons.

We identified L5 IB neurons in both V1 and vS1 based on their relatively large size and characteristic bursty firing pattern (Supplementary Figure S2). To mimic the feed-forward input, we injected a 2 ms current impulse (1.5 nA to 2 nA) to elicit spikes (Figure 5A and B). On some trials, paired with the feed-forward input, we delivered 2 ms 470 nm LED stimulation to activate feedback terminals. We chose an initial delay of 3 ms between the somatic current injection and subsequent optogenetic stimulation in order to maximize the effect on bursting based on previous reports ^40^. With feed-forward input only (Figure 5B left), there were low probabilities of bursting in both V1 and vS1 (Figure 5B, left). With both feed-forward and feedback inputs, we found a major difference in the response of neurons in V1 and vS1. For neurons in V1, LM feedback reliably elicited a single extra spike within 3 ms after LED stimulus onset, with a higher probability of second spikes (0.53 ± 0.15, mean ± s.e.m, same for the rest of the section) compared to the feed-forward-only condition (Figure 5A-C left, 0.20 ± 0.11, *p* = 0.045, Wilcoxon signed-rank test, *n* = 10 neurons). Subsequent spiking within at least a 20 ms time window was eliminated. This effect was consistent with the temporal sharpening effects we observed during sustained feed-forward excitation. In contrast, feedback from vM1 elicited bursts of spikes in in L5 vS1 IB neurons, with a higher probability of a second (0.60 ± 0.10) and a third spike (0.26 ± 0.06) compared to the feed-forward-only condition (Figure 5A-C; second spike with feed-forward only: 0.15 ± 0.07, *p* = 0.003, third spike with feed-forward only: 0.09 ± 0.04, *p* = 0.04, Wilcoxon signed-rank test, *n* = 13 neurons).

**Figure 5.**
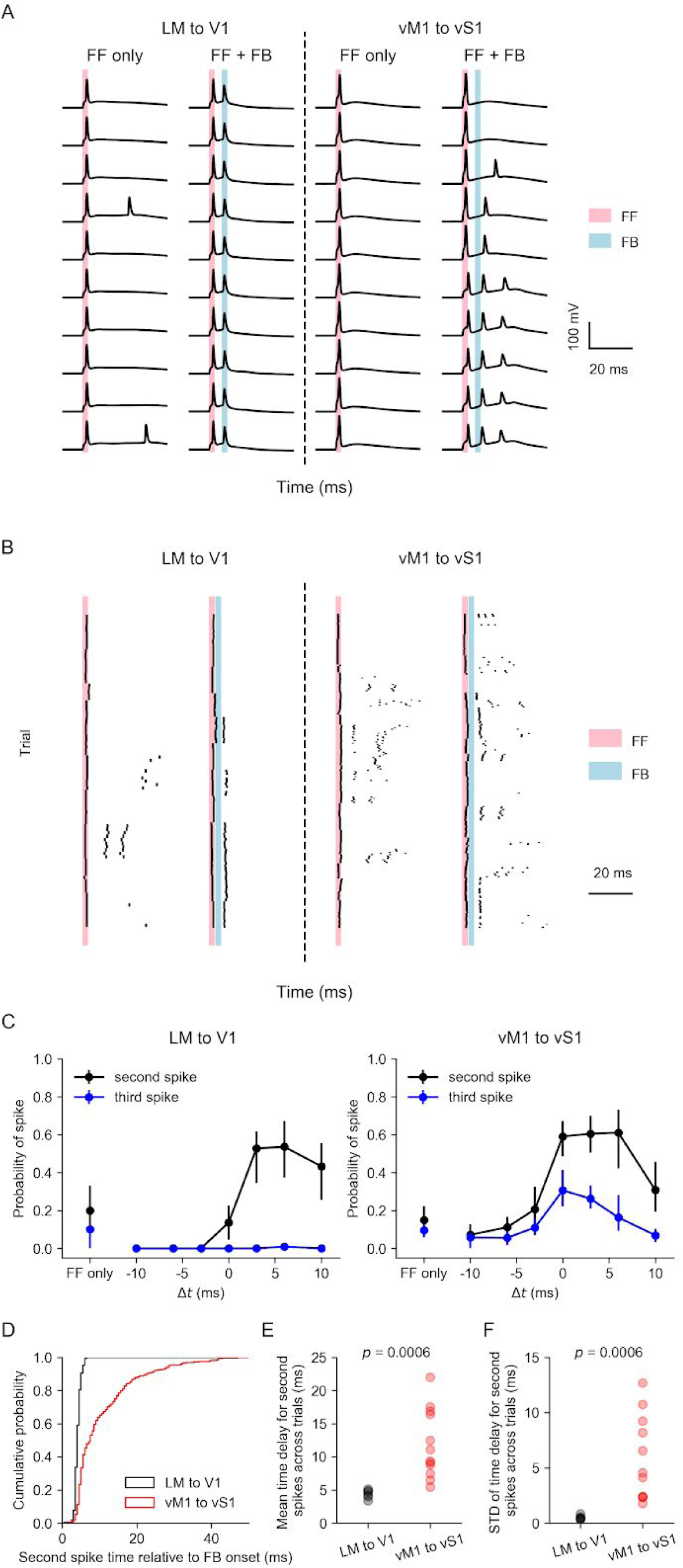
Feedback regulation on bursting behavior of L5 intrinsically bursty (IB) neurons. **(A)** Firing of example L5 IB neurons in response to feed-forward stimulus only (FF, 2 ms) and the combination of feed-forward and feedback stimulus (FF + FB, FB 2 ms) in V1 (left) or vS1 (right). In these examples, FB stimulus was delivered 3 ms after FF stimulus. **(B)** Raster plots of all cells in V1 (left) or vS1 (right). Each row refers to a trial and each tick indicates a spike. **(C)** Probability of occurrence of the second spike (black) and third spike (blue) of L5 IB neurons responsive to FF + FB stimulus, as a function of the time of FB onset relative to the FF onset (*Δt*). Dots and error bars are mean and bootstrapped 68% confidence intervals. **(D)** The cumulative probability of second spike occurrence time relative to the FB onset. Black: LM to V1. Red: vM1 to vS1 **(E) and (F)** Mean (E) and standard deviation (STD, F) of the time delay across trials for the second spikes were both significantly higher in the vM1 to vS1 pathway (*p =* 0.0006, Wilcoxon rank-sum test). Each dot represents a cell.

According to a previous study ^40^, the burstiness of L5 IB neurons is highly dependent on the time difference between the stimulation of the cell body (feed-forward) and the apical dendrites (feedback). We therefore varied the time delay (*Δt*) of the feedback onset relative to the feed-forward onset and characterized the corresponding probability of additional spikes contributing to bursting evoked by activating feedback. Feedback from LM to V1 only elicited an extra spike when the feedback was activated immediately after the feed-forward input, other than the *Δt* = 3 ms condition (Figure 5A and B left), the probability of second spikes at *Δt* = 6 ms (0.54 ± 0.14) was also significantly higher than the probability in the feed-forward-only condition (Figure 5C, left, *p* = 0.042, Wilcoxon signed-rank test, *n* = 10 neurons). Importantly, feedback from LM to V1 never elicited a third spike (Figure 5C, left). However, feedback from vM1 to vS1 increased the probabilities of both a second and a third spike when *Δt* was between 0 and 6 ms (Figure 5C, right). Regardless of *Δt*, vM1 to vS1 feedback never reduced the spike probability compared to the feed-forward-only condition.

Moreover, we also found that for the LM to V1 pathway, the timing of the second spike elicited by feedback was strongly locked to the feedback stimulus onset (Figure 5D), with a time delay of 4.4 ± 0.2 ms (mean ± s.e.m across neurons, *n* = 6 neurons), while the second spike timing in vM1 to vS1 pathway was both longer and more variable across trials (Figure 5D-F).

These results together suggest a major difference in the regulation of L5 IB cell bursting by the two feedback pathways: LM to V1 feedback induces a single, temporally-precise extra spike, similar as the behavior with sustained feed-forward input, while vM1 to vS1 feedback facilitates bursting of L5 IB neurons when feed-forward input was transient (2 ms).

## Discussion

In this study, we compared the wiring logic and functional consequences of feedback in two pathways: feedback between hierarchically organized cortical areas within a sensory modality (LM to V1) and feedback between motor and sensory areas (vM1 to vS1). We found major differences in the connectivity rules and the functional impact of feedback, arguing against a universal principle governing the organization and functional operation of feedback in the neocortex.

For the wiring, we found that feedback from vM1 to vS1 preferentially targeted VIP+ cells but formed much weaker connections to SOM+ cells. Our results confirmed the disinhibitory circuit in vM1 to vS1 feedback pathway ^21^, where feedback targets VIP cells who recruit SOM+ cells and disinhibit local pyramidal cells ^18^, the same circuit has also been found in the feedback from cingulate cortex to V1 ^24^. However, we found that this disinhibitory circuit was not preserved in the LM to V1 pathway. Instead, feedback from LM to V1 exhibited an opposite pattern with stronger connections to SOM+ cells compared to VIP+ cells, which, to our knowledge, is the first study that shows SOM+ cells are strongly activated by top-down feedback projections. Previous studies have revealed that SOM+ cells are involved in surround suppression because their activities increase with the stimulus size ^41,42^, and studies in both primates ^43,44^ and rodents ^45,46^ reveal that feedback from the higher visual area contributes to surround suppression. Our finding here filled in this gap by showing that feedback from LM to V1 recruits local SOM+ interneurons.

Functionally, we found that combined with sustained feed-forward input, both feedback projections temporally sharpened feed-forward excitation by eliciting a transient increase followed by a prolonged decrease in the firing rate of pyramidal cells. This “temporal sharpening” in our results is consistent with previous studies in a variety of contexts ^38,47–53^, suggesting that it is a canonical feature of neural circuits. The mechanism of the inhibitory phase of the biphasic response could include direct ^54^ or indirect ^55^ excitation of local interneurons from the long-range projections. Given the connectivity we described here, the most likely explanation of the inhibitory phase we observed is the direct excitation of interneurons from LM or vM1 feedback projections and not the indirect excitation of inhibitory cells via local V1 or vS1 excitatory neurons: despite we simultaneously drove a large number of feedback terminals with optogenetics, feedback excitation never elicited spiking in V1 or vS1.

Compared to the sustained feed-forward inputs, transient feed-forward inputs may better mimic the physiological conditions, because animals experience an external environment that changes rapidly. Here, we further showed that when paired with transient temporally-precise feed-forward input to the cell bodies, only feedback from vM1 to vS1 enhanced bursting responses in L5 IB cells, while feedback from LM to V1 continued to show a temporal sharpening effect. The difference in the connections to SOM+ cells and VIP+ cells between the two feedback pathways may play an important role in this process. VIP+ cells inhibit SOM+ cells that have been shown to target and gate the activity on the apical dendrites ^56–58^. Therefore, without the inhibition from SOM+ cells, such as in the vM1 to vS1 feedback pathway, the input on the apical dendrites from the feedback projections would induce plateau potentials, propagated to the cell bodies and elicit bursting ^40,59^. Previous work reveals that feedback from vM1 to vS1 elicits calcium spikes at the apical dendrites ^60^, and simultaneous activation of apical dendrites and cell bodies elicit bursting in L5 IB neurons ^40^. Here we linked these two findings together by directly showing that the combination of feed-forward and vM1 to vS1 feedback inputs were able to elicit bursting in L5 IB neurons in vS1. Along the previous arguments that bursts activate their targets stronger than single spikes, thus can prepare their targets for subsequent inputs ^61,62^, vM1 to vS1 feedback may also function as an attentional signal that amplifies the sensory responses. In the LM to V1 feedback pathway, in contrast, connectivity to SOM+ cells might explain why we did not observe bursting facilitation on V1 L5 IB cells, because of the inhibition from SOM+ cells to the apical dendrites. Other than SOM+ cells, we also found that feedback from LM to V1 targeted PV+ cells, consistent with previous reports ^23,63–65^. Different from SOM+ cells, the majority of PV+ cells target cell bodies instead. The fact that feedback from LM to V1 connects to both PV+ cells and SOM+ cells suggests that this feedback precisely controls the timing of activities not only at the cell body but also at the apical and tuft dendrites. Given the precise time window of the excitation, feedback from LM can only enhance the activity in V1 in cases where there is a temporal coincidence between higher-level representations in LM and sensory information in V1.

This leads to a question about what information V1 neurons integrate from LM feedback that requires such precise timing. Two major models that have been proposed for the feedback integration are the probabilistic inference model ^6^ and the predictive coding model ^4,5^. In both models, feedback is driven by a higher-order sensory feature, that could either be interpreted as prior information in the probabilistic inference model or the prediction signal in the predictive coding model. However, the two models give different predictions on how V1 neurons integrate feedback. The probabilistic inference model predicts that V1 neurons are excited by feedback encoding for similar features, which serves as a prior, while the predictive coding model predicts that V1 neurons are inhibited by feedback encoding for similar features, thus subtract the prediction and encode an error message. Both models require precise timing which ensures that only relevant activities are integrated. Since we found both excitation and inhibition on V1 principal cells, our finding is compatible with both hypotheses, depending on the specificity of the excitatory connections between LM and V1 principal neurons. Two recent studies ^37,45^ seem to provide evidence that LM to V1 feedback is more consistent with the predictive coding model ^66^. Marques et al. ^37^ reveals that LM to V1 feedback innervates locations in V1 that would be activated by stimuli orthogonal to or opposite to a cell’s own tuning, while Keller et al. ^45^ shows that V1 neurons are more likely to be innervated by the LM boutons whose receptive fields are offset from those of the V1 neurons. These results suggest that V1 neurons are precisely excited when feed-forward is driven by one feature (e.g. stimulus in a particular location) and feedback is driven by an orthogonal feature (e.g. stimulus in an offset location), encoding for an error message. In contrast, the activity of V1 neurons is suppressed if both feed-forward and feedback are driven by one feature (e.g. a uniform stimulus with large size), which is likely mediated by SOM+ cells, given the connectivity pattern we found and their size tuning properties ^41,45^. However, future work is needed to directly examine the structure-function connectivity rules between LM and V1 neurons.

## Acknowledgments

We thank Dr. Emmanouli Froudarakis, Dr. Dimitri Yatsenko, Dr. Saumil Patel, Yves Bernaerts and other lab members of Tolias lab for technical support. We thank Dr. Philipp Berens, Stelios Papadopoulos for discussions and comments on the manuscript.

## Author contributions

S. Shen, X. Jiang, F. Scala, and A. S. Tolias designed the experiments. S. Shen, X. Jiang, F. Scala, J. Fu, P. Fahey., and Z. Tan performed the experiments. S. Shen, X. Jiang, D.Kobak., and F. Sinz analyzed the data. S. Shen, X. Jiang, F. Scala, D. Kobak, F. Sinz, J. Reimer, and A. S. Tolias wrote the manuscript.

## Materials and methods

### Animals and surgeries

All procedures performed on animals were in accordance with the ethical guidelines of the National Institutes of Health and were approved by the Institutional Animal Care and Use Committee (IACUC) of Baylor College of Medicine.

In this study, we used 79 mice in total (51 males and 28 females), aged 8 weeks to 4 months. These included 4 C57Bl/6 mice (all male), 29 PV-Cre/Ai9 mice (19 males and 10 females), 35 SOM-Cre/Ai9 mice (22 males and 13 females), and 11 VIP-Cre/Ai9 mice (6 males and 5 females). All Cre and Ai9 reporter lines are on a C57Bl/6 background, and they are from Jackson Labs as follows:

SOM-Cre: https://jaxmice.jax.org/strain/013044.html

VIP-Cre: https://jaxmice.jax.org/strain/010908.html

PV-Cre: https://jaxmice.jax.org/strain/008069.html

Ai9 reporter: https://jaxmice.jax.org/strain/007909.html

Before each experiment, we performed the following surgical procedures on the animals. We used 3% isoflurane to induce anesthesia and anesthetized animals were placed in a stereotaxic head holder (Kopf Instrument). The anesthesia was then maintained with 1.5% to 2% isoflurane and the body temperature was maintained at 37°C during the whole surgical procedure using a 14 homeothermic blanket system (Harvard Instrument). We injected the following drugs at the beginning of the surgery: 0.05 mL, 0.5% bupivacaine subcutaneously under the scalp, 3 mg/kg dexamethasone intramuscularly in the leg, and 7.5 mg/kg ketoprofen subcutaneously on the back. After 10-20 minutes, we removed an approximately 1 cm^2^ area of skin above the skull and cleaned up the underlying fascia. Using the surgical glue (VetBond, 3M), we closed the wound margins and stuck the skins on the skull to avoid any open wounds. We then attached a custom-made head bar on the skull with dental cement (Dentsply Grip Cement). After the dental cement was completely dry, we removed the mouse from the stereotaxic frame and held the skull stationary on a small platform with the newly attached head bar. Using a surgical drill and HP 1/2 burr, we made a ∼3 mm craniotomy on the right hemisphere with a center 3 mm lateral of the midline and contacting the lambda suture on its posterior edge, which allowed the exposure of areas V1 and LM. The exposed cortex was then cleaned up with ACSF (125 mM NaCl, 5 mM KCl, 10 mM Glucose, 10 mM HEPES, 2 mM CaCl_2_, 2 mM MgSO_4_). After viral injections (will be described in the section *Virus Injections*), the cortical window was sealed with a 3 mm-diameter coverslip (Warner Instruments), using VetBond glue.

### Intrinsic optical imaging and visual areas identification

We used intrinsic imaging to identify the precise locations of V1 and LM (Figure 1B). The animal was kept anesthetized with 1-2% isoflurane. Similarly as previously described ^35^, we measured the change in cortex reflectance to red light with a wavelength of 610 nm. Using a CCD camera, we captured 512×512 pixels images at a rate of 12 Hz. To present visual stimuli, we positioned a 7’’ LCD monitor (Lilliput 665GL-70NP/HO/Y monitor, 60 Hz scan rate) approximately 10 cm away from the left eye of the animal, covering about 88° (azimuth) and 72° (elevation) of the contralateral visual field. To map the retinotopy of V1 and LM, we stimulated the animal with drifting gratings presented in one of the four locations (top lateral: -20° azimuth, 20° elevation; top medial: 20° azimuth, 20° elevation; bottom lateral: -20° azimuth, -20° elevation; bottom medial: 20° azimuth, 20° elevation) on each trial. The gratings were drifting either vertically or horizontally, with a spatial frequency of 0.03 cycles/°, a temporal frequency of 4 Hz, and a size of 8°. Stimuli were presented for 2 seconds and separated with a 3-second luminance-matched gray background. Stimulus displays were generated with MATLAB Psychophysics Toolbox and a photodiode attached to the screen that allowed a precise time-stamping of each frame of stimulus presentation on the clock of CCD camera recording. We then constructed the retinotopic map based on the brain reflectance using linear regression and identified V1, LM, and AL by comparing with the published retinotopic maps^31^.

### Virus injections

We used adeno-associated virus (AAV, serotype 2/1; University of Pennsylvania Gene Therapy Program Vector Core, titer 2.3×10^12^ /mL) to deliver the expression of ChR2-EYFP, under the CaMKIIα promoter, which allows ChR2-EYFP to be only expressed in excitatory neurons in LM or vM1.

Anatomical structures used in this study V1, LM, vS1, vM1, are shown in Figure 1A. To inject in LM, we made the injection in one of the four retinotopic locations of LM, with the guidance of the vessels and the retinotopic map obtained with intrinsic imaging. To inject in vM1, we stereotactically injected with coordinates 1.1 mm anterior and 0.9 mm lateral relative to bregma (Figure 1A). Virus was injected into two depths in the cortex, 300 μm and 700 μm below pia, with a volume of 30 nL each, aiming to cover cells in all layers.

Two to four weeks after the injections, we checked the virus infection by imaging the EYFP expression pattern under a two-photon microscope (Figure 1C). We took two-photon 300 μm stacks, covering the whole craniotomy.

### Multi-cell whole patch-clamp recording in brain slices with optogenetics

The visual cortical slice preparation generally followed the protocol in the previous study ^56^, with the addition of recently developed NMDG (N-Methyl-D-glucamine) adult animal slicing method ^67^.

Before the experiments, we injected 2% Fast Green FCF (Sigma-Aldrich) stereotactically into the area in V1 that had the densest expression of ChR2-EYFP. Then the animal was put into deep anesthesia with 3% isoflurane and decapitated. The brain was quickly removed and placed into 0-4 °C oxygenated NMDG solution (93 mM NMDG, 93mM HCl, 2.5 mM KCl, 1.2 mM NaH_2_PO_4_, 30 mM NaHCO_3_, 20 Mm HEPES, 25 mM glucose, 5 mM sodium ascorbate, 2 mM Thiourea, 3 mM sodium pyruvate, 10 mM MgSO_4_ and 0.5 mM CaCl_2_, pH 7.35). We cut 300 μm thick parasagittal slices from the tissue blocks with a microslicer (Leica VT 1200). With the guidance of the injected dye, we only kept the slices that contain our region of interest in V1 or vS1 (LM or vM1 was not in the slices), which was marked with Fast Green. Slices were kept at 37.0 ± 0.5 °C in oxygenated NMDG solution for 10-15 min and then transferred to the normal ACSF (125 mM NaCl, 2.5 mM KCl, 1.25 nM NaH_2_PO_4_, 25 mM NaHCO_3_, 1 mM MgCl_2_, 25 mM glucose and 2 mM CaCl_2_, pH 7.4) for 0.5-1 h before recording. During the recording, the slices were submerged in a chamber and stabilized with a fine nylon net attached to a platinum ring. The recording chamber was perfused with the oxygenated ACSF.

We performed multi-cell whole-cell patch recordings in L2/3, L4 and L5 in the region of interest in V1 or vS1, with 4-7 MΩ borosilicate pipettes (2.0 mm OD, 1.16 mm ID, Sutter Instruments) filled with a standard low-chloride internal solution (120 mM potassium gluconate, 10 mM HEPES, 4 mM KCl, 4 mM MgATP, 0.3 mM Na_3_GTP, 10 mM sodium phosphocreatine and 0.5% biocytin, pH 7.25). For IPSC recordings, a Cs+ based internal solution was used instead (135 mM cesium methanesulfonate, 10 mM HEPES, 2.5 mM MgCl_2_, 4 mM Na_2_ATP, 0.4 mM Na_3_GTP, 10 mM sodium phosphocreatine, 0.5 mM EGTA, 0.1 mM spermine and 0.5% biocytin, pH 7.25). On each slice, we always recorded at least one L2/3 pyramidal cell as a reference. For voltage-clamp recordings, we clamped the voltage at -85 mV to record EPSC. For current-clamp recordings, we adjusted the membrane voltage to -70 mV (Figure 2B-E). To activate ChR2 expressed in the axon terminals, we delivered 470 nm LED blue light through the light path of the microscope. 2 ms LED pulses were used to trigger EPSCs or EPSPs (Figures 2 and 3). In some experiments to identify monosynaptic events (Figure S3), we applied TTX (1 μM) to block action potentials, and then applied 4-AP (0.5 mM) to block potassium channels and thus enhance the responsiveness ChR2 expressing axon terminals. If the EPSC was recovered with 4-AP, we regarded it as a monosynaptic event.

Cell classes were primarily identified by their genetic markers and were further confirmed with electrophysiological and morphological features (Figure S1 and S2). Several recent studies have shown that SOM-Cre lines also label some fast-spiking (FS) cells ^68–70^. FS cells have fundamental differences in the morphology and function from other SOM+ types ^71^ (Figure S1). We, therefore, excluded FS cells in the SOM+ group in later analyses.

In experiments for temporal sharpening (Figure 4), we injected 200-800 pA positive current lasting 300 ms to drive the cells to fire with a regular pattern. On LED-on trials, we delivered 20 ms 470 nm LED stimuli in a random time, ranging from 100 ms to 200 ms after the onset of the current injection. LED-off trials were interleaved with the LED-on trials, to control for the effect by the firing pattern change across time.

In experiments for bursting (Figure 5), we identified the L5 intrinsic bursty neurons by their laminar position, large cell bodies, and thick apical dendrites. After patching the cells, we then confirm the burstiness with their firing pattern (Figure S2). We injected 2 ms 2000 pA positive current to elicit a single spike. The intrinsic bursty neurons sometimes fire a burst of spikes in response to the current injection only. On LED-on trials, we delivered 2 ms 470 nm photostimuli with various time differences relative to the current injection: -10, -6, -3, 0, 3, 6, 10 ms. For each time difference, we recorded 6 to 10 trials.

### Data analysis for EPSC or EPSP recordings

#### Amplitude and latency measurement

The baseline activity was defined as the mean activity within 40 ms prior to the LED stimulus onset. The peak value of EPSC or EPSP was the maximum value within 35 ms after the LED stimulus onset relative to the baseline activity. If the cell fired in response to the light stimulus, we then defined the EPSP amplitude as the difference between the baseline membrane potential and threshold potential. The latency of the synaptic event was estimated from the extrapolated intersection of the baseline with a line through the two points of time when the current was 20% and 80% of the peak value ^72,73^.

#### Criteria of cell “responsiveness”

We considered a cell responsive to light stimulation (Figure 3A) when both criteria were met: first, the latency of the response (EPSC or EPSP) is less than 8 ms; second, the amplitude of the response is larger than 3 times the standard deviation of the baseline activity.

#### Statistical comparison of EPSC or EPSP values

To compare the amplitudes of EPSCs and EPSPs across different slices and animals, we normalized the amplitude of individual cells to the amplitude of the L2/3 pyramidal cell on the same slice. If there were more than one L2/3 pyramidal cells recorded on the same slice, we normalized by their mean amplitudes.

We statistically compare the normalized amplitudes of EPSC or EPSP between different cell classes across slices within the same feedback pathway. We performed Conover’s test ^74^ for comparisons among cell classes other than L2/3 pyramidal cells, which is the *post hoc* pairwise comparison followed by Kruskal-Wallis one-way analysis of variance by ranks. We performed a permutation signed test for comparisons between L2/3 pyramidal cells and other types.

To statistically compare the normalized amplitudes of EPSC or EPSP of the same cell type across different pathways, we performed the Wilcoxon rank-sum test, with *p* values corrected by Benjamini-Hochberg adjustments for multiple comparisons.

#### Criteria for “spiking cells”

To quantify the probability of neurons to fire in response to light stimulation (Figure 3C), we only included data from slices with a minimal level of ChR2-expressing feedback innervation, defined by a minimum mean EPSC in L2/3 pyramidal cells of 50 pA. We never observed spikes on slices that did not meet this minimum criterion.

### Data analysis of experiments with a sustained current step

For the experiments with sustained current steps (Figure 4), we detected the spikes by thresholding and aligned the spike trains with the onset of LED pulses. We computed mean firing rates across trials for each cell, which were smoothed by convolving with a box-car filter of 4 ms time bins. To quantify the excitatory effect of the feedback modulation, we compared the firing rate of LED-on trials and LED-off trials within 10 ms after the LED stimulus onset of the LED-on trials or the matching time points of the LED off trials. To quantify the inhibitory effect of the modulation, we compared the spike delay of the first spike after the LED-on period of the LED-on trials or the matching LED-on period of the LED-off trials, relative to the LED stimulus onset. We performed the Wilcoxon signed-rank test for the statistical comparison.

### Data analysis of bursting experiments

For the bursting experiments (Figure 5), we performed two analyses. First, we showed the probability of spiking of the second spike and third spike as a function of the time difference between the feed-forward and feedback inputs (Figure 5B). We performed the Wilcoxon signed-rank test to compare the probability of second spikes and third spikes with feedback stimulus delivered at 3 ms after feed-forward onset to those with feed-forward stimulus only. Second, for those trials that a second spike occurred, we characterized the time delay of the second spike relative to the feedback onset. For each cell that had at least a second spike across trials, we computed the mean and standard deviation of the time delay, and we compared these two values of cells in the two feedback pathways with Wilcoxon rank-sum test.

## Supplementary Materials

### Confirmation of cell classes with their recovered morphologies and firing patterns

In a subset of experiments, we recovered the morphologies of the recorded cells to confirm their cell classes. Morphologies and firing patterns were particularly important in two cases in our experiments. First, it has been reported in previous studies that a subset of SOM+ cells are fast-spiking (FS) basket cells (Figure S1A and B, bottom), while the rest of the cells are non-fast-spiking (Non-FS) cells (Figure S1A and B, top). As the two subsets of cells are very different in the morphology and firing pattern (Figure S1A and B), we excluded the activities of FS cells in the main analyses, because their behavior is closer to PV+ cells (Figure S1C and D). Second, in the bursting experiments (Figure 5), we mainly used the firing pattern to identify the intrinsic bursty cells in L5 in V1 or vS1. The firing pattern of the busty cells was featured with their high firing rate (> 100Hz, Figure S2) at the beginning of the current injection and the large afterhyperpolarization (AHP) following the bursting. We morphologically recovered a subset of the cells with bursty firing patterns and consistent with previous literature, these cells were thick-tufted cells (Figure S2).

**Figure S1.**
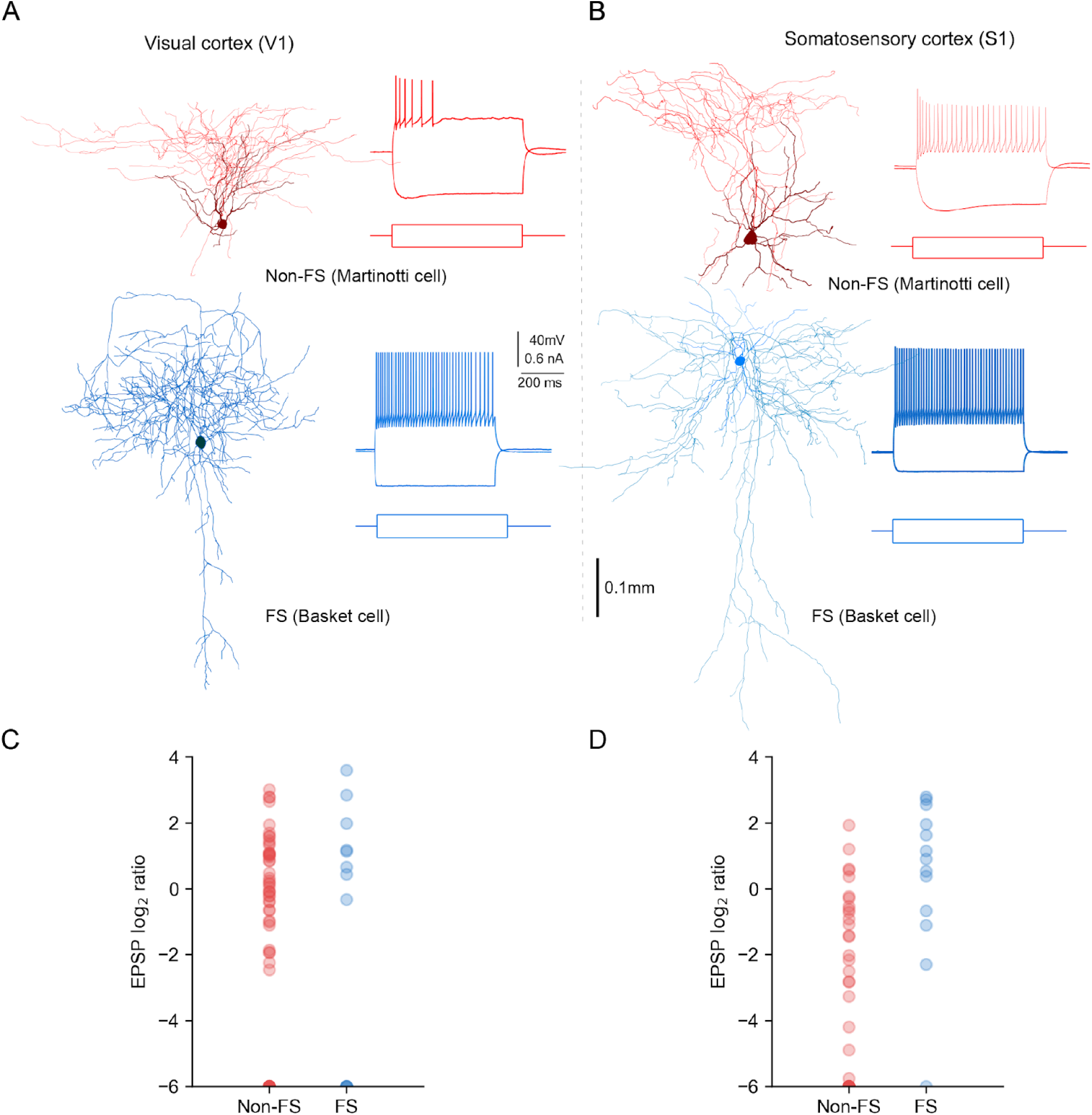
SOM+ cells are composed of cells with different firing patterns and morphologies. **(A)** Non-fast-spiking (Non-FS) cells and fast-spiking (FS) cells of SOM+ cells in V1. Example morphologies and firing patterns of Non-FS cells (top, red) and FS cells (bottom, blue). In this example, the Non-FS cell was a Martinotti cell and the FS cell was a Basket cell. **(B)** FS cells and Non-FS cells of SOM+ cells in vS1. Example morphologies and firing patterns of Non-FS cells (top, red) and FS cells (bottom, blue). In this example, the Non-FS cell was a Martinotti cell and the FS cell is a Basket cell. **(C)** Log normalized EPSP of Non-FS cells and FS cells in V1. **(D)** Log normalized EPSP of Non-FS cells and FS cells in vS1.

**Figure S2.**
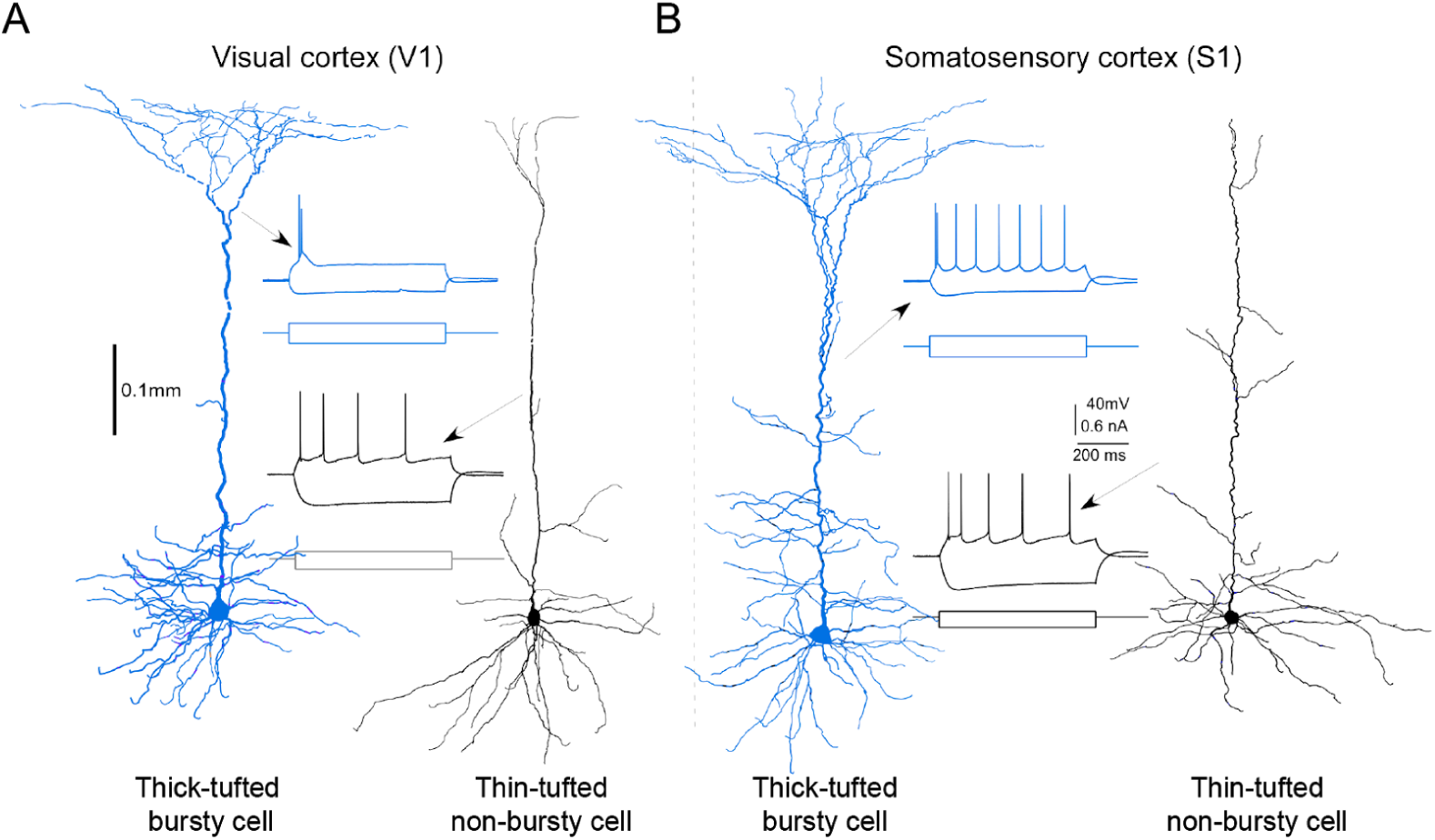
Intrinsic bursty cells and Non-bursty cells in L5 were identified with both the morphologies and the firing patterns. **(A)** Morphologies and firing patterns of example thick-tufted bursty cell (left, blue) and thin-tufted non-bursty cell (right, black) in V1. **(B)** Morphologies and firing patterns of example thick-tufted bursty cell (left, blue) and thin-tufted non-bursty cell (right, black) in vS1.

### Most of the recorded EPSC and EPSP events were monosynaptic

To confirm that the elicited events were monosynaptic, in a subset of the connectivity experiments, we applied the sodium channel blocker tetrodotoxin (TTX, 1 µM) and the potassium channel blocker 4-aminopyridine (4-AP, 500 µM) ^1^. TTX blocks action potentials to prevent polysynaptic events, while 4-AP enhances ChR2-mediated depolarization of axonal boutons, increasing the detectability of monosynaptic events. We found that all light-evoked synaptic events were silenced by TTX alone, but most of them could be partially recovered by the addition of 4-AP (Figure S3A and B), indicating that most of the excitatory responses we recorded were indeed monosynaptic. The latencies of most of the evoked events (3/83) recovered with the addition of 4-AP were less than 8 ms (Figure S3C). Thus we used 8 ms as a threshold to identify monosynaptic events in all subsequent experiments. It is possible that some events by this standard were polysynaptic, but we found that the recorded excitatory cells were never driven to fire by feedback activation (Figure 3E), indicating a low rate of polysynaptic events.

**Figure S3.**
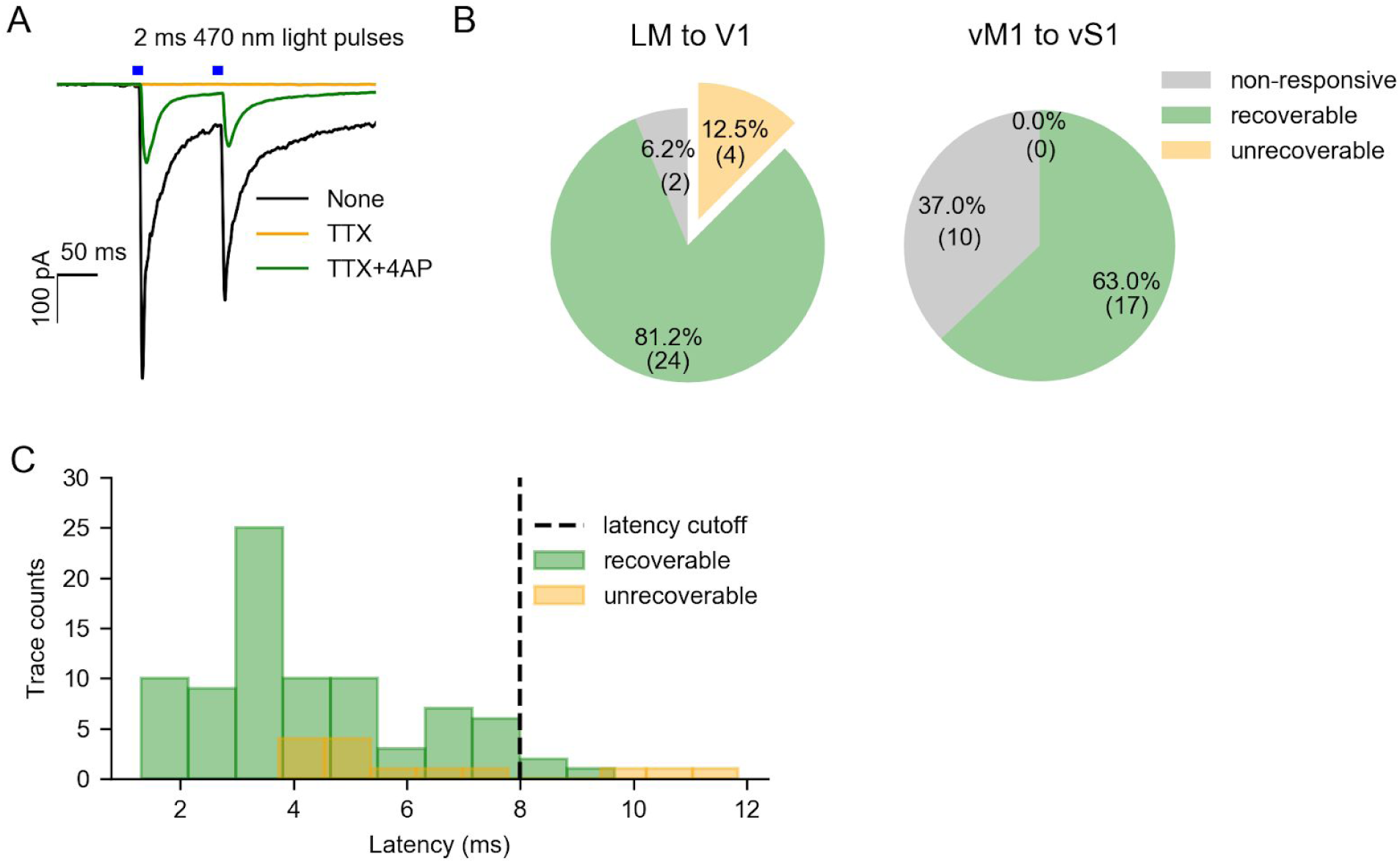
Identify monosynaptic events with pharmacological methods. **(A)** Example traces of feedback (blue bars) responses with no drug (black), TTX (orange), and both TTX and 4AP (green). The cells that were still responsive in the presence of TTX and 4AP were identified as recoverable cells. **(B)** Proportion of cells that are non-responsive without drug (gray), recoverable (green) and unrecoverable (orange) in V1 (left) or vS1 (right). **(C)** Latency distribution of recoverable traces (green) and unrecoverable traces (orange). We set 8 ms as the latency cut-off for “responsive” cells in the rest of the experiments.

**Table S1.**
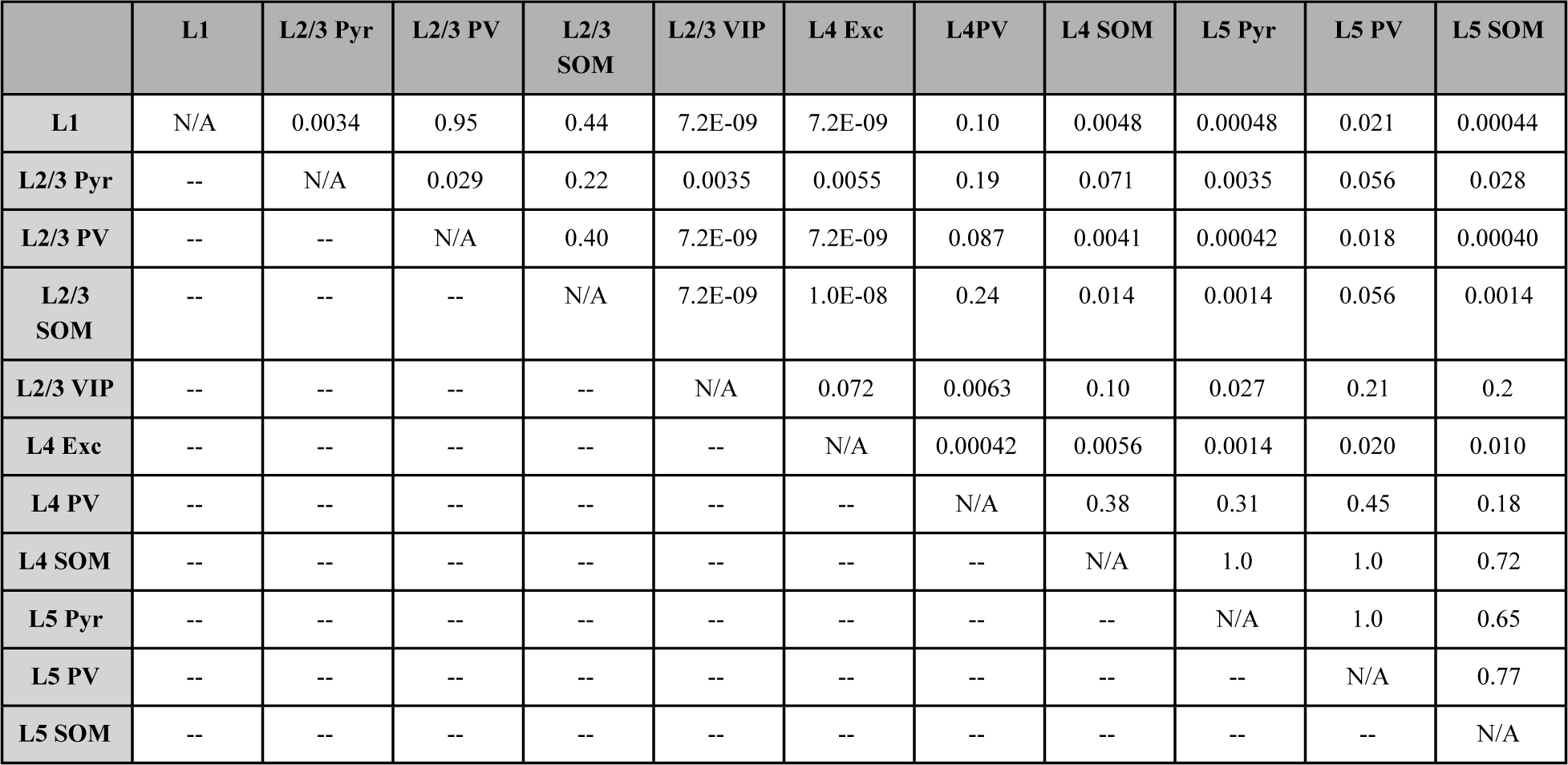
*p* values of LM feedback elicited normalized EPSC comparison across cell classes in V1. Conover’s test for comparisons among cell classes other than the L2/3 pyramidal cells. Permutation signed test for comparisons between L2/3 pyramidal cells and other cell classes.

**Table S2.**
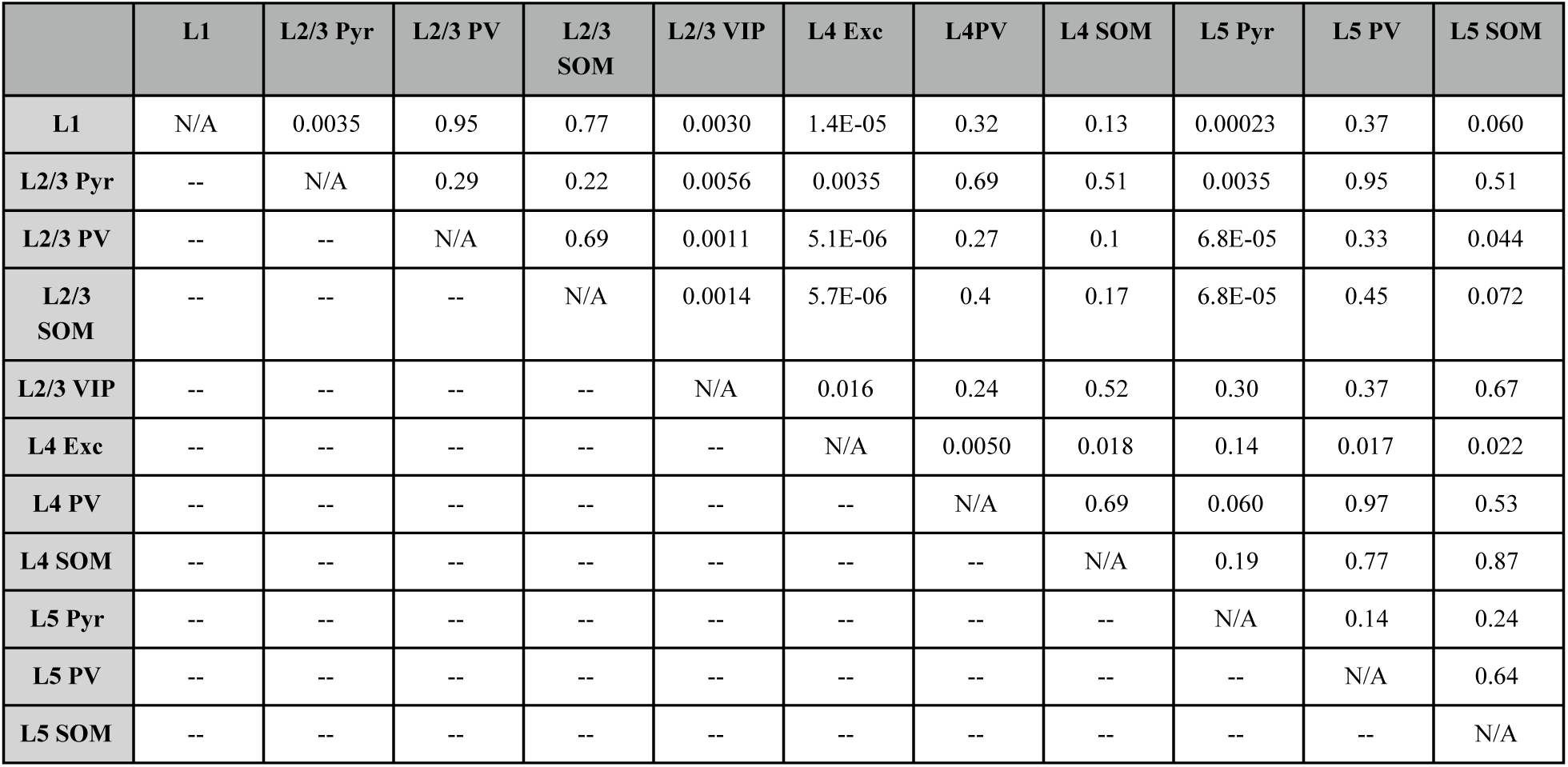
*p* values of LM feedback elicited normalized EPSP comparison across cell classes in V1. Conover’s test for comparisons among cell classes other than the L2/3 pyramidal cells. Permutation signed test for comparisons between L2/3 pyramidal cells and other cell classes.

**Table S3.**
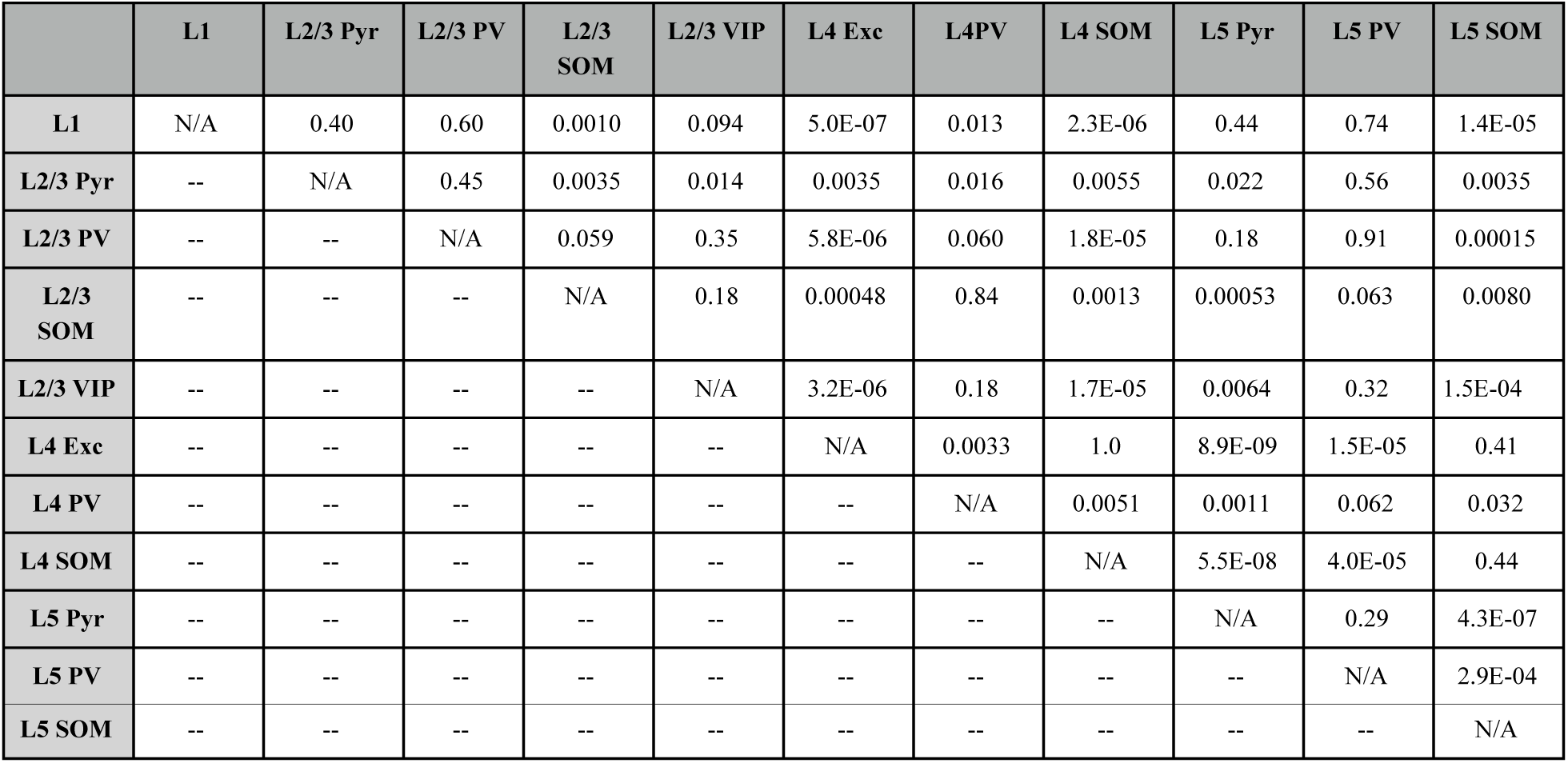
*p* values of vM1 feedback elicited normalized EPSC comparison across cell classes in vS1. Conover’s test for comparisons among cell classes other than the L2/3 pyramidal cells. Permutation signed test for comparisons between L2/3 pyramidal cells and other cell classes.

**Table S4.** *p* values of vM1 feedback elicited normalized EPSP comparison across cell classes in vS1. Conover’s test for comparisons among cell classes other than the L2/3 pyramidal cells. Permutation signed test for comparisons between L2/3 pyramidal cells and other cell classes.

|  | L1 | L2/3 Pyr | L2/3 PV | L2/3 SOM | L2/3 VIP | L4 Exc | L4PV | L4 SOM | L5 Pyr | L5 PV | L5 SOM |
| --- | --- | --- | --- | --- | --- | --- | --- | --- | --- | --- | --- |
| L1 | N/A | 0.032 | 0.19 | 0.041 | 0.64 | 0.0001 | 0.0051 | 0.00012 | 0.75 | 0.25 | 0.00016 |
| L2/3 Pyr | -- | N/A | 0.31 | 0.048 | 0.038 | 0.014 | 0.014 | 0.032 | 0.17 | 0.35 | 0.0056 |
| L2/3 PV | -- | -- | N/A | 0.62 | 0.21 | 0.0043 | 0.17 | 0.0056 | 0.35 | 0.81 | 0.0087 |
| L2/3 SOM | -- | -- | -- | N/A | 0.0068 | 0.0038 | 0.27 | 0.0056 | 0.1 | 0.37 | 0.0087 |
| L2/3 VIP | -- | -- | -- | -- | N/A | 1.7E-06 | 0.00034 | 5.2E-06 | 1 | 0.27 | 3.5E-06 |
| L4 Exc | -- | -- | -- | -- | -- | N/A | 0.059 | 1 | 0.00027 | 0.00092 | 0.74 |
| L4 PV | -- | -- | -- | -- | -- | -- | N/A | 0.073 | 0.015 | 0.062 | 0.13 |
| L4 SOM | -- | -- | -- | -- | -- | -- | -- | N/A | 0.00043 | 0.0014 | 0.75 |
| L5 Pyr | -- | -- | -- | -- | -- | -- | -- | -- | N/A | 0.44 | 0.00056 |
| L5 PV | -- | -- | -- | -- | -- | -- | -- | -- | -- | N/A | 0.0021 |
| L5 SOM | -- | -- | -- | -- | -- | -- | -- | -- | -- | -- | N/A |

### Raw values of EPSC and EPSP show that vM1 to vS1 feedback pathway might have an overall weaker connections than LM to V1 pathway

In the main text (Figure 3), we have normalized the EPSC and EPSP amplitudes to those of L2/3 pyramidal cells to account for the variability in the amount of virus injection across the animals. In Figure S4, we show the raw values of EPSC and EPSP. Under the assumption that the strength and amount of virus injected are similar among animals, we found that EPSC and EPSP of most of the cell classes in the vM1 to vS1 pathway are significantly lower than those in LM to V1 (Figure S4, Table S5), suggesting an overall weaker connection of vM1 to vS1 feedback pathway. Note that the raw EPSC values in VIP+ cells were similar between V1 and vS1, but raw EPSP values were higher in vS1 than in V1 (*p* = 0.0067, Wilcoxon-rank test, *p* values adjusted with Benjamin-Hochberg method). This is because the input resistance of VIP+ cells in vS1 (median 287 MΩ) is higher than that of VIP+ cells in V1 (median 204 MΩ, Figure S5, *p* = 1.8E-05, Wilcoxon rank-sum test). With a similar amount of synaptic current, VIP+ cells in vS1 responded with a higher EPSP.

**Figure S4.**
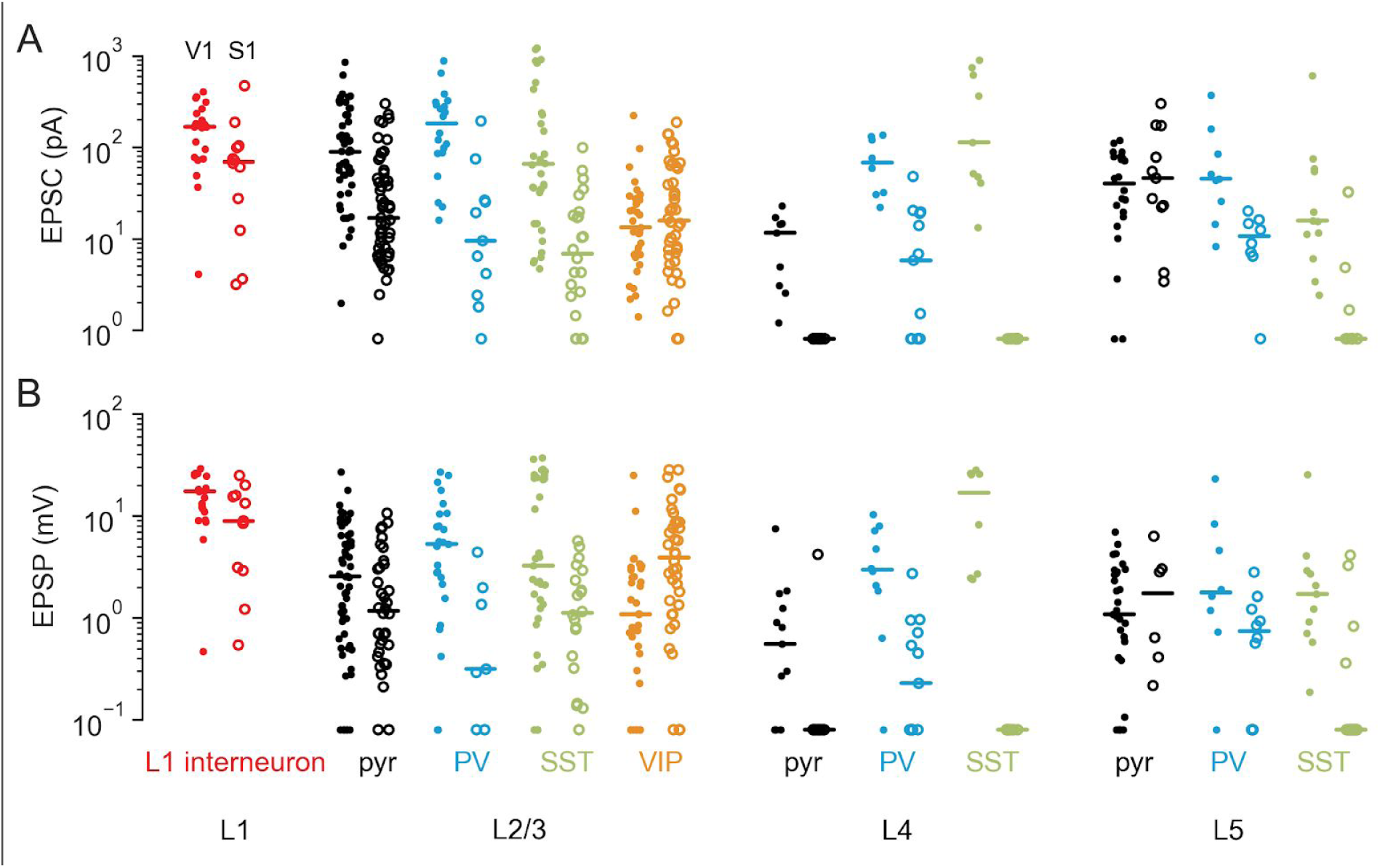
raw EPSC (A) and raw EPSP (B) of different cell classes in response to the feedback activation.

**Figure S5.**
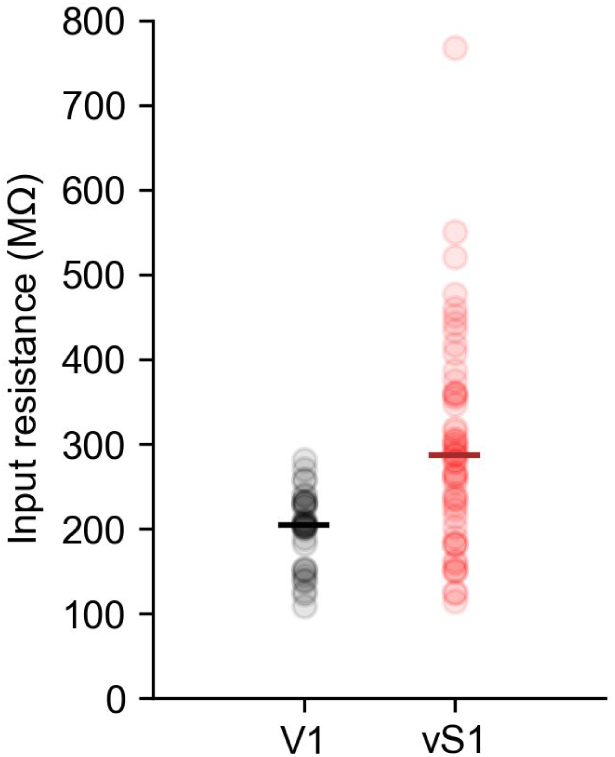
Input resistance of VIP+ cells in V1 or vS1. Each dot represents a cell and the bar marks the median of the population. VIP+ cells in vS1 have significantly higher input resistance than those in vS1 (*p* = 1.8E-05, Wilcoxon rank-sum test).

**Tables S5.**
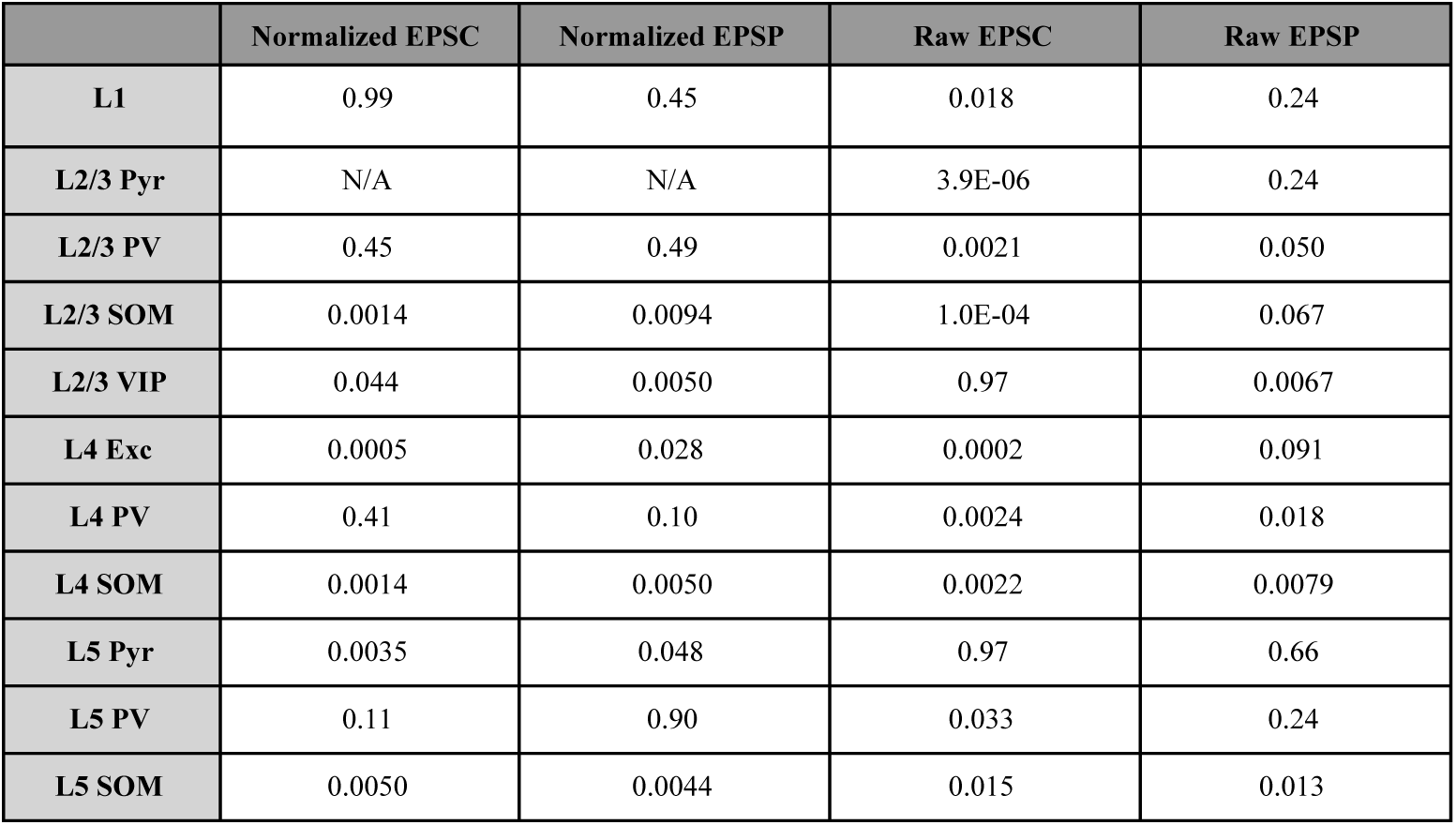
*p* values of comparison of normalized EPSC and EPSP of the same cell type across the two feedback pathways. Wilcoxon-rank test, *p* values adjusted with Benjamin-Hochberg method.

|  | Normalized EPSC | Normalized EPSP | Raw EPSC | Raw EPSP |
| --- | --- | --- | --- | --- |
| **L1** | 0.99 | 0.45 | 0.018 | 0.24 |
| **L2/3 Pyr** | N/A | N/A | 3.9E-06 | 0.24 |
| **L2/3 PV** | 0.45 | 0.49 | 0.0021 | 0.050 |
| **L2/3 SOM** | 0.0014 | 0.0094 | 1.0E-04 | 0.067 |
| **L2/3 VIP** | 0.044 | 0.0050 | 0.97 | 0.0067 |
| **L4 Exc** | 0.0005 | 0.028 | 0.0002 | 0.091 |
| **L4 PV** | 0.41 | 0.10 | 0.0024 | 0.018 |
| **L4 SOM** | 0.0014 | 0.0050 | 0.0022 | 0.0079 |
| **L5 Pyr** | 0.0035 | 0.048 | 0.97 | 0.66 |
| **L5 PV** | 0.11 | 0.90 | 0.033 | 0.24 |
| **L5 SOM** | 0.0050 | 0.0044 | 0.015 | 0.013 |

### Variability of “temporal sharpening” effect across cells

In Figure S6 and S7, we show more example cells on their sharpening effect. While the “temporal sharpening” effect of L2/3 and L5 pyramidal cells were highly consistent across cells (Figure S6 and Figure S7, A and C), the effect on excitatory cells in L4 was variable (Figure S6 and Figure S7B).

**Figure S6.**
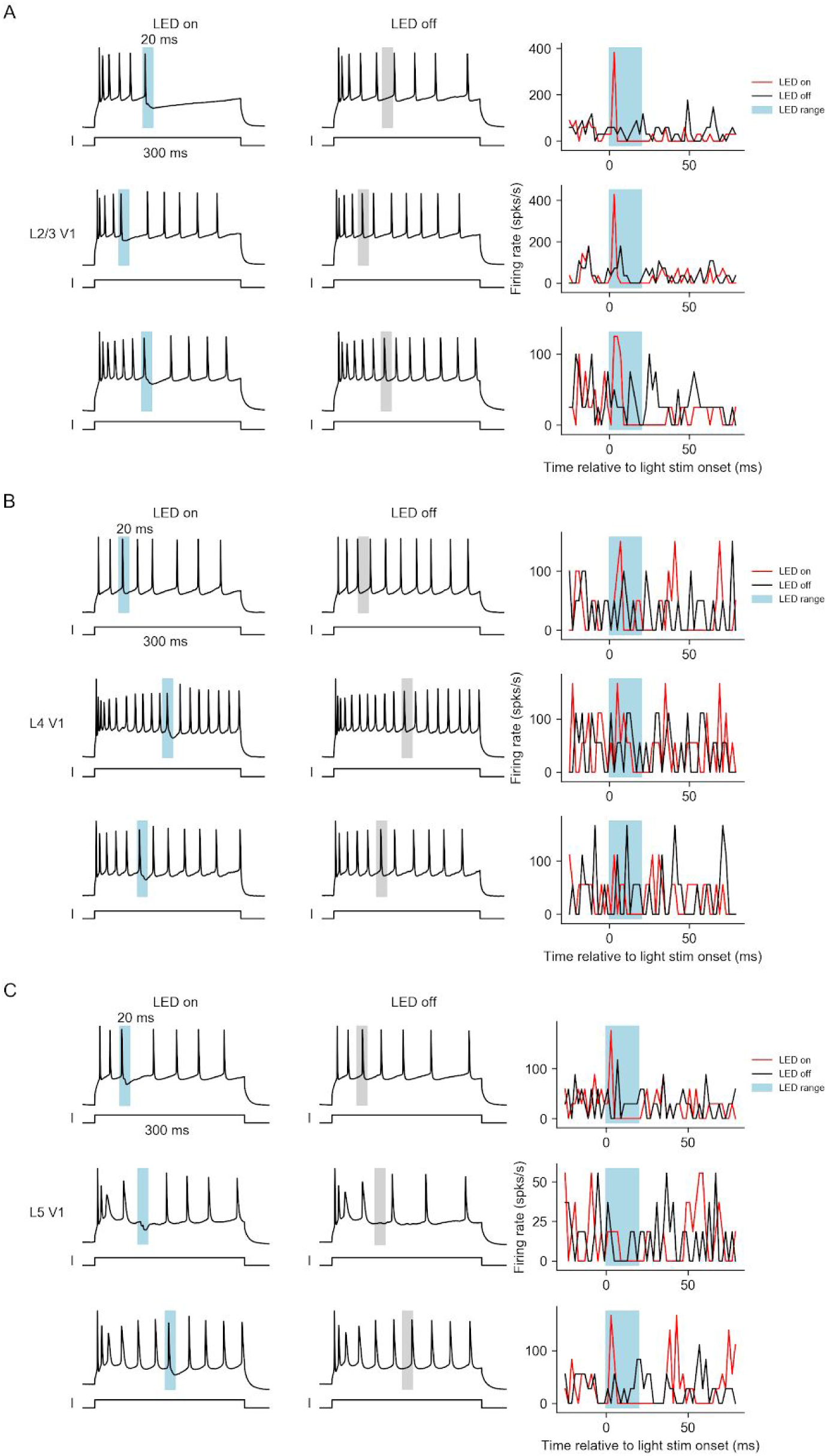
More example V1 principal cells in L2/3 (A), L4 (B), or L5 (C) with sustained feed-forward current and brief feedback inputs. Left and Middle: example traces with LED on (left) or LED off (middle).

**Figure S7.**
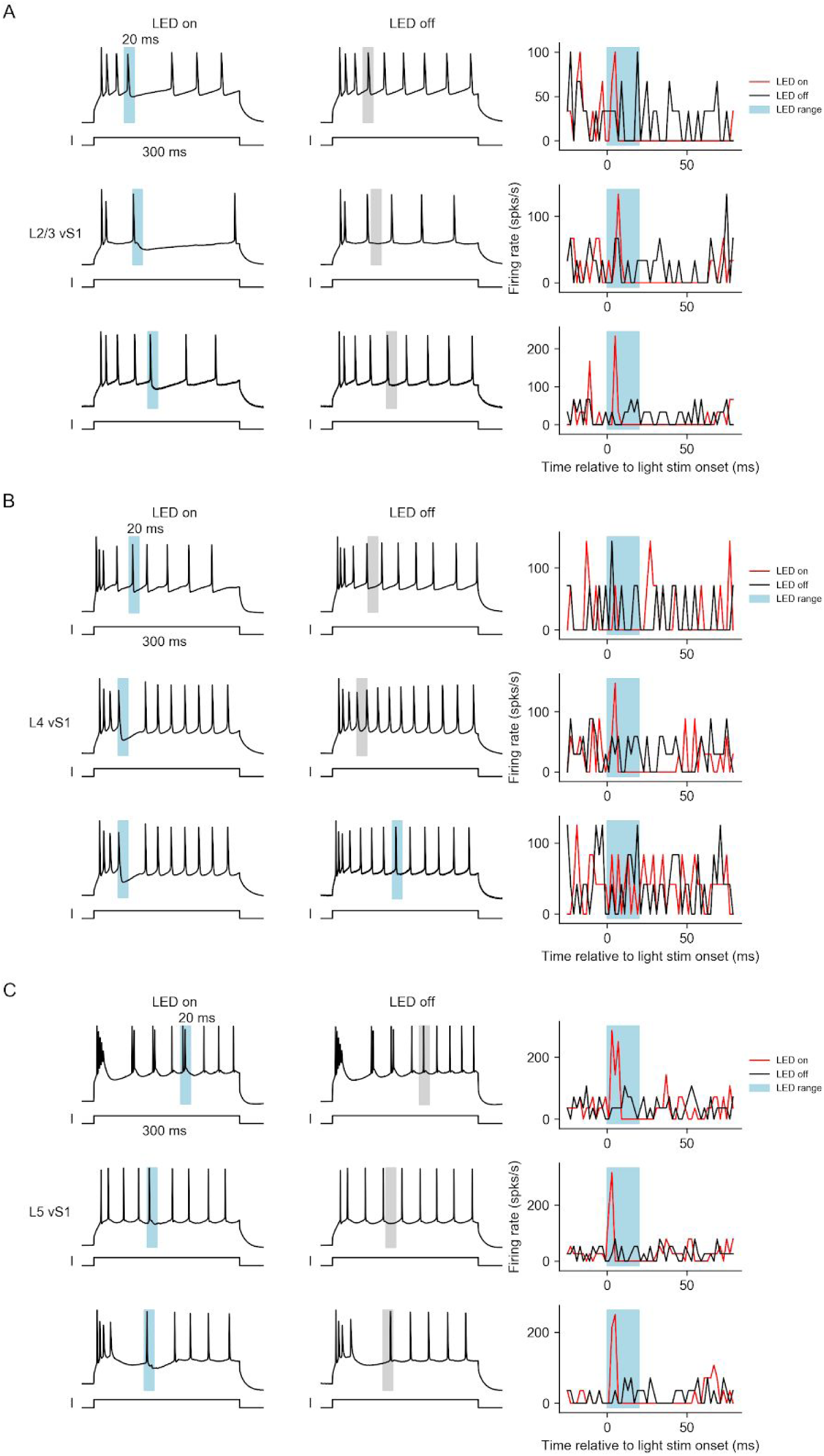
More example vS1 principle cells in L2/3 (A), L4 (B), or L5 (C) with sustained feed-forward current and brief feedback inputs. Left and Middle: example traces with LED on (left) or LED off (middle).

